# Analysis of hospital air microbiomes uncovers potential avenues for transmission of diverse pathogens linked to hospital-acquired infections

**DOI:** 10.64898/2026.03.13.711524

**Authors:** Julio Cesar Ortega Cambara, Oscar Previtali, Piotr Cuber, Pedro Humberto Lebre, Shanom Ali, Darren Chooneea, Raju Misra, Hermine V Mkrtchyan

**Affiliations:** Centre for Innovation in Genomics and Microbiome Sciences, School of Medicine and Biological Sciences, University of West London, London, United Kigdom; Centre for Microbial Ecology and Genomics, University of Pretoria, Pretoria, South Africa; University College London Hospital NHS Trust, London, United Kingdom; UK Health Security Agency, London, United Kingdom

**Keywords:** Low-biomass microbiome, ONT long read sequencing, metagenomics, antimicrobial resistance, hospital-acquired infections

## Abstract

Fomite-mediated and airborne transmission pathways play a significant role in the dissemination of healthcare associated pathogens, contributing to the burden of hospital-acquired infections (HAIs). Effective infection prevention and control therefore require robust surveillance approaches capable of capturing the complexity of air and surface microbial communities present in healthcare settings. However, to date such approaches are not widely employed. In this study we used a rapid and low-biomass optimized metagenomic workflow to detect clinically relevant pathogens, characterise genetic signatures such as antimicrobial resistance and virulence determinants, and profile the wider airborne microbiome, including unculturable taxa essential for HAI surveillance.

Air samples were collected from multiple clinical and non-clinical areas across three multi-storey healthcare units within a UK hospital, alongside to surface samples from a haematology/oncology ward with a documented history of outbreaks. Following DNA extraction and enrichment optimised for low-biomass samples, sequencing was performed using the Oxford Nanopore Technologies MinION long-read platform. Metagenomics data were analysed using an in-house developed bioinformatics pipeline.

Metagenomic profiling revealed high bacterial taxonomic diversity across sampled environments, with limited overlap between the airborne and surface communities. Approximately 0.3% of bacterial reads harboured a large variety of antimicrobial resistance genes (ARGs), virulence factors (VFs) and mobile genetics elements (MGEs) across different hospital sample groups. Notably, critical and emerging pathogens were detected across ten wards and were associated with clinically significant resistance determinants, including multiple *bla_OXA_* subtypes, *vanA* gene clusters, and multidrug efflux pumps conferring resistance to eight important classes of antibiotics, including carbapenems, cephalosporins, penams, and vancomycin. In addition, several plasmid replicons involved in horizontal gene transfer (HGT) were identified within these pathogens, indicating an increased potential for the emergence and dissemination of multi-drug resistance within hospital associated microbial communities.

Our findings highlight the importance of using rapid metagenomics-based methodologies for environmental surveillance in healthcare settings. Correspondingly, this study revealed that hospital air and surface microbiomes comprise complex and dynamic microbial communities that harbour divers**e** of ARGs and VFs with the potential for rapid transmission across the surface-air interface, particularly in high-risk/vulnerable patient areas where outbreaks are more likely to occur.

## 1. Background

Hospital associated infections (HAI) continue to challenge healthcare systems worldwide. More than 3.5 million cases of HAI are estimated to occur in the EU/EEA each year, resulting in over 90,000 deaths, a burden that exceeds the cumulative burden of other infections^1^. Antimicrobial resistant (AMR) bacteria, including bacteria resistant to last-resort antibiotics, such as carbapenem-resistant Enterobacterales, are responsible for 71% of HAIs cases in the EU/EEA alone^1^, and AMR infections are also associated with increased patient hospitalisation and mortality^2^. AMR-important pathogens may spread easily, either directly from person to person or indirectly between the environment and people^4^. Environmental sources have been linked with respiratory disease, whereby transmission of infectious disease and antimicrobial resistance genes (ARGs) occurs through inhalation^4^. Around 300,000 people per year, in England alone, acquire HAIs during National Health Services (NHS) care^5^, which can be transmitted through contact with surfaces (including sinks, drains, ventilation systems and frequently touched surfaces)^6^ and via airborne particulate matter and dust^7^. Most hospital wards are poorly ventilated, and the windows are sealed with too few single rooms. Such environments not only increase the risk of infection transmission but also provide a unique opportunity for the spread of AMR by horizontal gene transfer (HGT)^8^. Hospitals therefore may become both a hub for transmission and a reservoir for antibiotic resistance, and such larger sources of antibiotic pollution can create new AMRs^9,10,11^. Additionally, existing infection control practices may have little or no effect on pathogens and AMR spreading through the air and surfaces widely^12,13^.

The rise of antimicrobial resistance (AMR) is a global crisis which threatens the lifesaving role of antibiotics, modern medicine is built upon. Clinical environments, including air and surfaces, harbour diverse microbes such as viruses, bacteria, fungi, and other eukaryotes, acting as potential reservoirs for pathogen and AMR transmission^14,15^. Microorganisms, including nosocomial pathogens, can survive on dry surfaces for up to several months, depending on factors like surface material and environmental conditions^16^. Swab tests revealed that Gram-positive bacteria, such as *Clostridium difficile*, can persist from days to seven months, while Gram-negative bacteria like *Klebsiella spp.* survive from hours to 30 months^17^. Viruses can last from a few days (e.g., SARS, Influenza)^18^ to several months (e.g., astrovirus, Herpes)^16,19^.

Despite the critical role of environmental reservoirs, including surfaces and air in the transmission of infectious diseases and AMR, these reservoirs have historically been poorly characterised due to technological limitations, including insufficient sensitivity, high costs, and slow processing times^20,21^. Most diagnostic testing and genomic surveillance strategies rely on expert identification using modern molecular and phenotypic techniques e.g. PCR, MALDI-TOF MS, phenotypic assays and targeted (gene or single species) next-generation sequencing^22^. However, each of these approaches suffers the same problems, they are targeted and therefore biased to a specific species, often culture-based and therefore time-consuming. Consequently, they fail to detect a substantial proportion of environmental microorganisms, particularly those present at low concentrations, or those that cannot be easily culturable in laboratory conditions^23^. These limitations reveal a critical gap in strategies to mitigate healthcare-associated infections (HAIs), AMR, and their dissemination via environmental reservoirs.

Although this gap has been emphasised repeatedly in the literature, recent technological advancements are beginning to address it. These include the use of metagenomic studies to investigate the diversity transmission of AMR across microbiomes in the healthcare setting^24,25,26^. Nevertheless, significant challenges remain, particularly within hospital environments. Previous studies have investigated air and surface microbiomes in healthcare settings have taken targeted approaches, with some focusing on specific pathogens and others adopting a broader taxonomic approach^27,28,29^. These studies revealed the intricate dynamics of complex microbial communities and functional genomic traits within healthcare environments and the potential of air and surface microbiome profiling as a powerful tool to inform infection control strategies and reduce the spread of HAIs and AMR.

We hypothesise that leveraging recent technological advances to develop pipelines for improved sampling, DNA sequencing and data analysis for low-biomass microbiomes (hospital air samples) can be adapted to more deeply characterise pathogens linking AMRs with species identification from hospital environments. This will, in turn, will support hospital sampling and analytical strategies for outbreak prevention and HAIs control.

In this study we have compared air and surface sampling and characterisation methods within a hospital setting, using compact, portable air collectors coupled with traditional surface sampling approaches. Together these sampling methods were paired with Nanopore driven long-read, portable, shotgun metagenomic sequencing. Our in-house developed bioinformatics pipeline enabled the detailed determination of both the taxonomic composition of the air and surface microbiome, their functional annotations, including antibiotic resistance traits, virulence factors and mobile genetics elements. This rapid and efficient pipeline demonstrated the use of air and surface surveillance of microbial diversity, detection, and characterisation of pathogens in hospital settings. It enables the detection of pathogens, their genomics signatures as well as characterisation of the wider microbiome, including those that are unculturable and often are missed by traditional testing methods, critical for the control of HAIs and tackling the global crisis of AMR.

## 2. Materials and methods

### 2.1. Clinical setting and sample collection

Areas were selected at random from a multi-storey healthcare teaching hospital trust in London consisting of 700-bed multi-specialty wards. The hospital hosted low risk patient groups, pre-assessment clinics and high-risk/vulnerable patients requiring augmented care (haematology, elderly care, adolescent haematology/oncology and infectious diseases). Each ward was a single floor of the hospital building assessed and included both outpatient and inpatient settings. Shared-occupancy wards included bed bays (room with 4-6 beds) with joint-access to bathrooms while patients residing in single-isolation rooms (SIRs) had sole access to a dedicated en-suite bathroom.

A total of 25 air and 36 surface samples were collected over five weeks period between March - April 2021 from both clinical and non-clinical areas including: Emergency Department (ED), Intensive Care Unit (ICU), Infectious diseases (ID) ward, Adolescent haematology/oncology, Oncology (Adult), Haematology (Adult), Elderly Care, Radiology, Surgical Ward, Theatres and Staff-only areas (**Table S1; Figure S1**). All surface samples were taken from a single ward (ward E), while air samples were taken from 10 different wards, with a minimum of two samples per ward.

**Figure S1:**
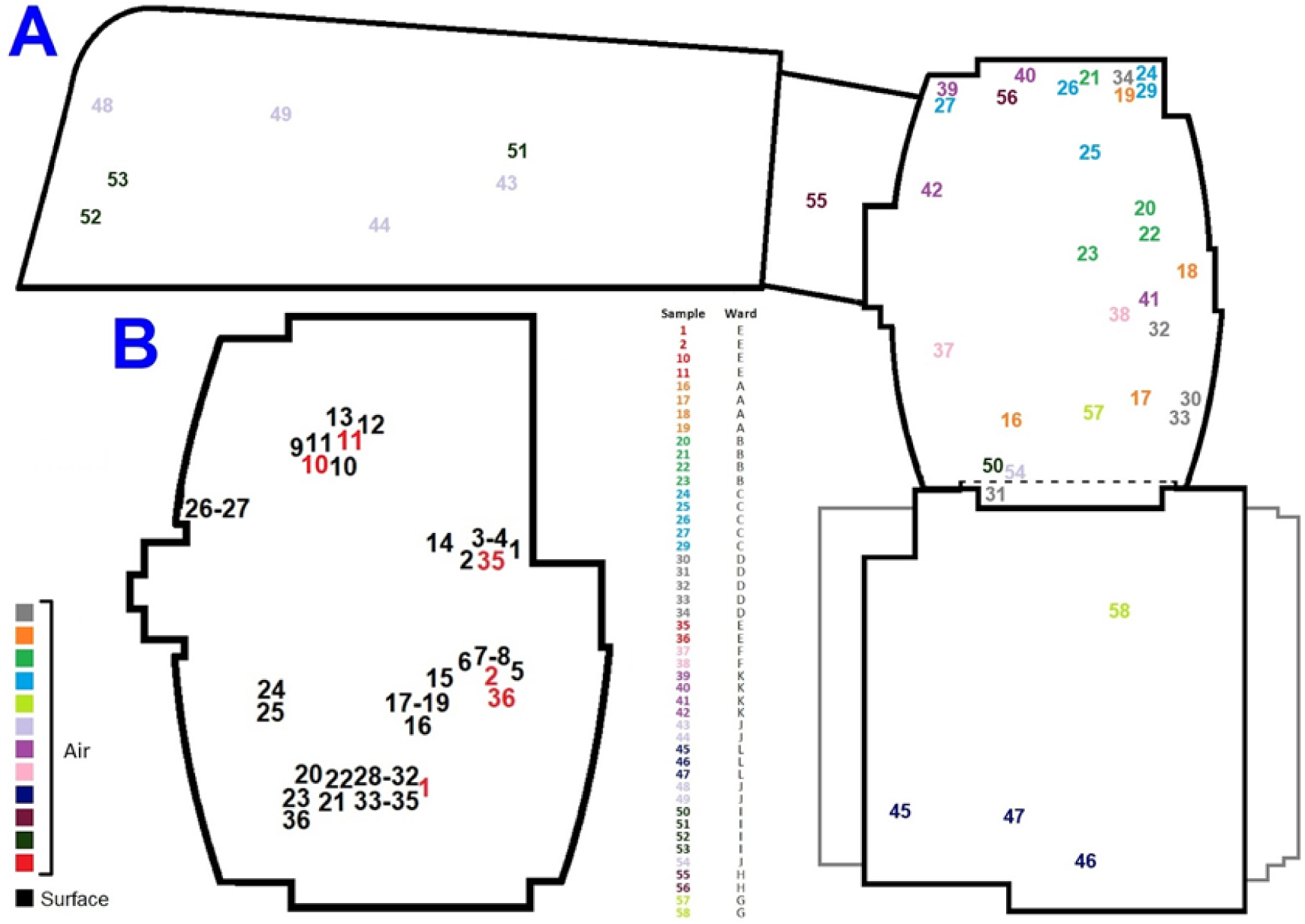
**A**- distribution of collection sites for both air and surface samples in the hospital: different colours indicate different wards (different floors from top to bottom) as well as surface vs air samples. **B** - Details of the ward where both air and surface samples were taken. Legend: **Air samples** - Kitchen: 1, 22; Private bed: 2, 20, 21, 24, 29, 31, 35, 36, 39, 40, 52; Clean Room (CR52): 11; Dirty room: 25; Staff only area: 16, 42, 45; Corridor: 17, 51, 53, 57, 58; Bed bay: 19, 26, 27, 30, 33, 34, 37, 41, 50; Reception: 23, 38; Treatment room: 32, 43, 44, 48, 49, 54, 55, 56; Decontamination unit: 46; Laboratory: 47. **Surface samples** - Vent: 1, 2, 5, 6, 9, 10, 12, 14-16, 20, 21, 28-32; Fridge: 3, 4, 7, 8, 17-19, 24, 25, 33-35; Fridge: 3, 4, 7, 8, 17-19, 24, 25, 33-35; Macerator side port: 11; Smoke detector: 13; Freezer: 22, 23, 26, 27, 36.

### 2.2. Air sample collection, transportation, and processing

Air samples were collected using a portable and battery powered Coriolis air sampler (Coriolis Micro, Bertin Instruments, France) for 1 hour per sample. For each sample, 3 ml was collected at a flow rate of 300 Lt/min resulting in 18 m^3^ volume of air collected. For this study, we used Invitrogen™ UltraPure™ DNase/RNase-Free water (Life Technologies Ltd., Paisley, UK) that was dispensed into the collection cone prior to sample collection. Collection cones were sterilised by autoclaving before each use. Approximately 3 ml of collected sample was then placed into 5 mL sterile Eppendorf tubes immediately after collection, on site. Samples were transferred to the labs at UWL within 1 hour of collection and stored at -80°C until further processing. Air sample DNA extraction and enrichment were carried out using a previously patented protocol (patent number: EP4562140A1).

### 2.3. Surface Swab sample collection, transportation and processing

All surfaces were sampled and processed following methods previously described [86]. Briefly, surfaces were wiped (in a left-to-right motion, followed by wiping at 45° and 90° angles; the process was repeated 3 times) using a 25 cm^2^ sponge swab (TSC Technical Service Consultants Ltd, UK) premoistened with neutralising solution A (0.1% [wt/vol] sodium thiosulfate). Sponge samples were transferred to refrigeration (2-8°C) within 2 hours of collection and processed within 24 hours.

To quench the activity of residual biocide on surfaces sampled, sponge swabs were further neutralised by placing them aseptically into sterile stomacher bags (VWR International, United Kingdom) containing 20mL neutralising solution B (which comprised: 3% (w/v) Tween 80 (Sigma-Aldrich, UK), 0.3% (w/v) Lecithin (Sigma-Aldrich, UK), 2% (w/v) Sodium thiosulfate (Sigma-Aldrich, UK), 1.5% (w/v) K2HPO4 (VWR Chemicals, UK), 0.5% (w/v) KH2PO4 (Sigma-Aldrich, UK), 1% Poly-[Sodium-4-Stryrenesulfonate] (Sigma-Aldrich, UK), 0.1% (w/v) Triton x100 (Sigma-Aldrich, UK) and prepared in Phosphate-buffered saline (PBS). Neutraliser solutions were sterilised by autoclaving at 121°C for 15 minutes and refrigerated at 2-8°C until required. The contents of each stomacher bag were homogenised manually by vigorously massaging the bag between the fingertips for 1 minute and incubated at ambient temperature for 10 min. Aliquots (500 µL) of the homogeneous solution were transferred to 1.5 mL Eppendorfs and stored at -80°C until further processing.

Sequential elution from a 500 µL aliquote was performed with a 0.45 µm membrane (Advantec MFS, Inc., Dublin, California) followed by a 0.22 µm Durapore® Hydrophilic PVDF membrane (Cat. No. GVHP01300, Millipore, Billerica, MA) and a 0.1µm PVDF membrane placed in a swinny filter using a luer-lock 10ml syringe. Samples then underwent DNA extraction using the same protocol specified above for the air samples, and quantified using Qubit dsDNA HS.

### 2.4. Metagenomics Sequencing of air and surface samples

DNA samples were sequenced following the ONT’s Rapid Barcoding (SQK-RBK004) protocol (Oxford Nanopore Technologies, UK) with adjustments to add volumes of final reagents, (SQB = 56 µL, LB = 7 µL, nuclease free water = 0 µL). These ratios were found to be optimized for low input samples. The libraries were loaded unto R9.4.1. flow cells (Oxford Nanopore Technologies, UK), either MinION (FLO-MIN106D) or Flongle (FLO-FLG001). Base-calling was performed using the MinKNOW® workflow, with a minimum Q score threshold of 10. All runs generated > 1000 FASTQ reads per file using a super-accurate basecalling model.

### 2.5. Bioinformatics pipeline

#### Prefiltering and taxonomic classification

Only reads classified as “pass” reads by the MinKNOW® platform were used for subsequent bioinformatics analysis. To filter out human contamination, reads were mapped against the human genome (GRCh38.p14) using Minimap2 v2.24^30^, and the resulting unmapped reads were used for taxonomy profiling. For reproducibility, taxonomy profiling was processed using the nf-core/taxprofiler bioinformatics pipeline v1.1.8 (https://github.com/nf-core/taxprofiler)^31,32,33^. Briefly, FastQC v 0.12.1 was used for read quality control, porechop v 0.2.4 for adapter clipping and merging (https://github.com/rrwick/Porechop), Filtlong v0.2.1 for low complexity and quality filtering (https://github.com/rrwick/Filtlong), and Minimap2 v2.24 for host-read removal. Taxonomic classification and profiling was done using Kraken 2 v2.1.2^34^ and the NCBI database. The contaminants were analysed using Recentrifudge^35^, which implements a robust method for the removal of the negative controls and crossover taxa from the rest of the samples. Additionally, a script was built to remove the potential contaminants from the sample groups affected.

#### AMR, VFs and MGEs detection

Identification of antimicrobial resistance, virulence genes, and mobile genetics elements, was performed on the classified reads using ABRicate v1.0.1 (https://github.com/tseemann/abricate) with the CARD^36^, VFDB^37^ and PlasmidFinder databases^38^. The fastq files were converted to fasta files using seqkit v2.8.2^39^. The ARG/VFs-carrying reads were used for the subsequent analysis of the co-existence of ARGs, VFs and MGEs. The pathogens were identified according to the published WHO priority list^40^.

#### Statistical analysis and visualization

All statistical analyses and visualisation were performed in Rstudio with the R (v4.1.3) software^41^. Results with p-values and adjusted p-values (q-values) < 0.05 were considered as statistically significant. Differences in the bacterial microbiome composition between air and surface samples were calculated using the Bray-Curtis index on the species-level counts extracted from the Kraken2 outputs, which were normalized using relative abundance percentages. The resulting matrix was visualized using a Non-metric Multidimensional Scaling (NMDS) plot, and the statistical significance of the compositional differences between air and surface sample groups was estimated using a PERMANOVA test. The differences in genetic elements (ARGs, VFs and MGEs) composition between different sample groups were estimated using the binary Jaccard index on a matrix containing information regarding the presence/absence of genetic elements per taxa per sample and visualized using a Principal Coordinate Analysis (PCoA) plot. PERMANOVA was also used to estimate the significance of the differences between air and surface samples according to presence/absence of genetic elements. All diversity metrics and statistical tests were calculated using the vegan package in Rstudio^42^. The Microbiome Multivariable Associations with Linear Models (MaAsLin 3) package^43^ were used to estimate the differential abundance of genetic element classes between the two sample groups. Counts for classes of genetic elements were generated by summing the counts for the genes in each specific class and then normalised to TPM by dividing them by the “per million” scaling factor of each sample (number bacterial reads/1,000,000).

Exploration of the hospital microbiome and the genetic makeup was conducted using a R shiny application called “hosMicro”, developed as part of this research (https://github.com/Julio92-C/hosMicro/). The Venn diagrams were graphed by “VennDiagram” package, the heatmaps were visualized with the “pheatmap” package^44^, the chord diagram was visualized with “circlize” package^45^ in R. The other figures were plotted by “ggplot2” and coloured with “paletteer” packages^46^.

A multi!zllayer bipartite network was constructed to visualise associations between air!zlsurface microbiomes and genetic elements (ARGs, VFs, MGEs). Samples were linked to detected taxa using abundance!zlweighted edges, while taxa were connected to their associated genetic elements using binary edges. The network was imported into Gephi (v0.10)^47^ and spatialised with the Fruchterman–Reingold algorithm to reveal clustering patterns. Node types were colour!zlcoded, and network metrics (degree centrality, betweenness, modularity) were calculated to identify key taxa and genetic elements.

## 3. Results

### 3.1 Microbial composition of the air and surface microbiome

The Kraken2 classification of the sequenced reads resulted in a total of 2636227 reads classified as bacterial, representing 36 phyla, 1627 genera, and 6509 species. Samples across the dataset shared the majority (6336; 97.3%) of bacterial species, with 86 (1.3%) and 87 (1.3) being unique to air and surface samples, respectively (**Figure S2A**). However, the significant clustering (PERMANOVA *R*^2^ = 0.117; *p-value* < 0.009) of these sample groups into distinct clusters according to their species composition (**Figure S2B**) suggested that the unique species in each group have a disproportionate importance to the community composition of the air and surface microbiomes. Indeed, this was observed in the distribution of the top 10 genera for each group (**Figure 1**), which revealed that air samples were dominated by *Rahnella* (37.6 % average rel. abundance) and *Paracoccus* (10.6% average rel. abundance), while surface samples were dominated by *Pseudomonas* (31.2% average rel. abundance) and *Psychrobacter* (14.4% average rel. abundance). It is important to note that despite these general trends, sample microbial composition was highly variable, particularly for surface samples.

**Figure 1.**
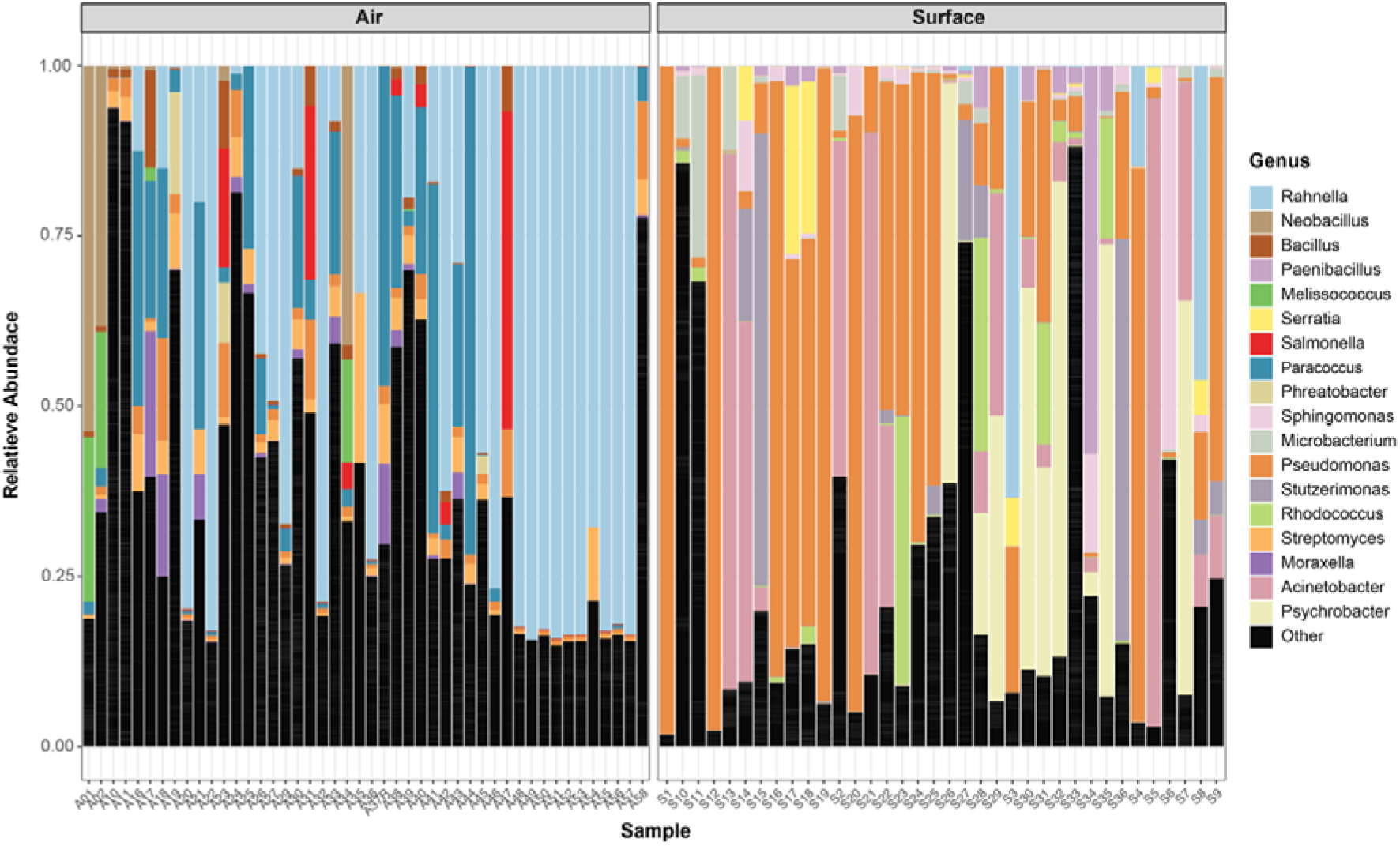
Relative abundance of the top 10 genera per sample group (air vs surface), based on the average relative abundance of genera across each group. Genera not identified as the top 10 are represented by the black bars as “Other”.

**Figure S2.**
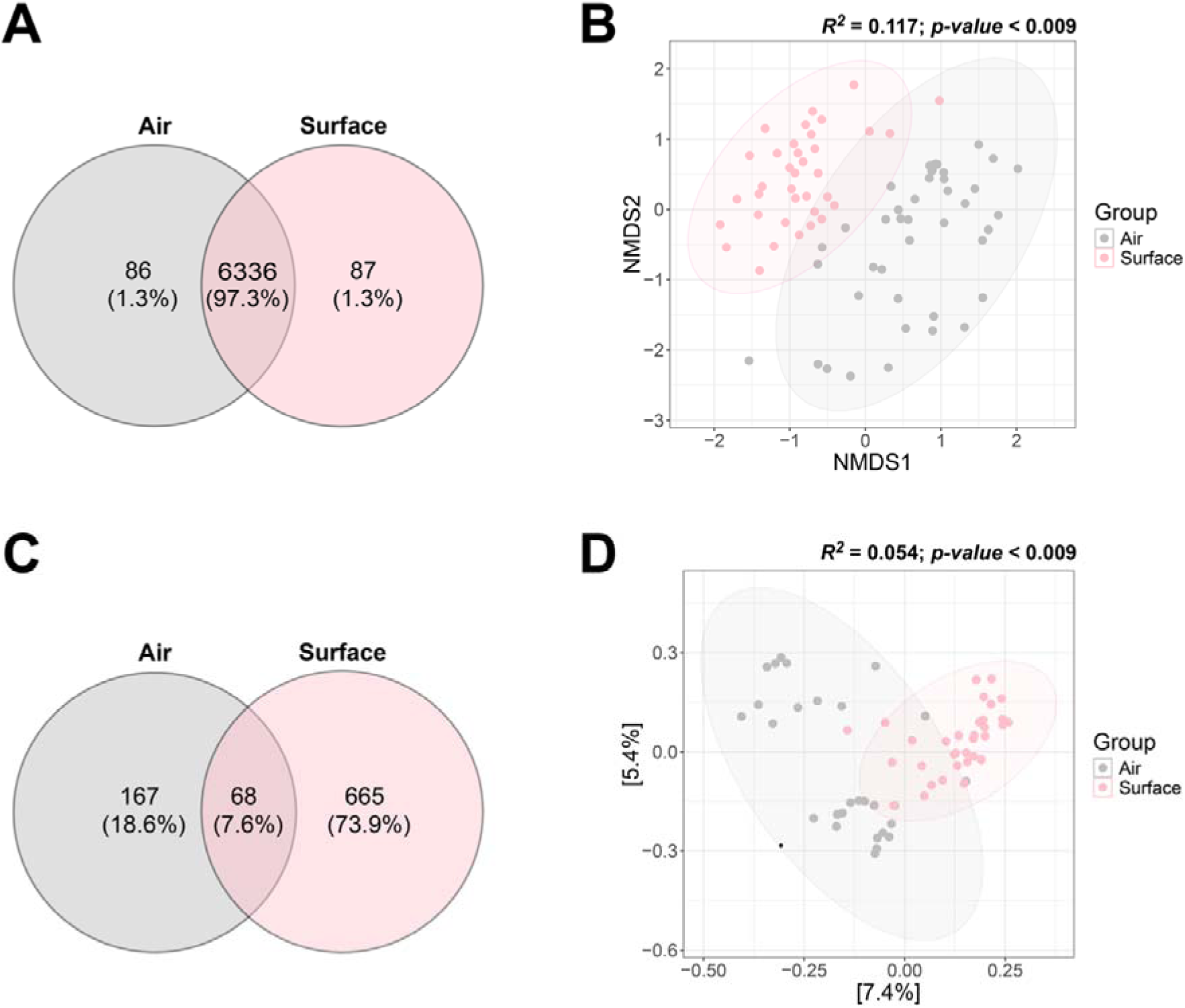
Diversity of the microbiome in air and surface samples according to kraken2-classified reads and contigs containing relevant genetic elements. Venn diagrams represent the number of bacterial reads classified at the species level (**A**) and contigs with genetic elements (**C**) unique to air and surface samples, as well as those shared by both groups. The ordination plots display the Bray-Curtis dissimilarity matrix between samples according to the abundance of bacterial species in the dataset (**B**), as well as the binary Jaccard dissimilarity matrix according to the presence/absence of contigs with relevant genetic elements (**D**). The PERMANOVA tests for the clustering of samples according to sample groups (air vs surface) are displayed in the legends for each ordination plot.

Genetic elements associated with AMR, VFs, and MGEs were detected in a total of 8122 contigs across 25 air samples and 33 surface samples. These were further classified into 900 taxa across all seven taxonomy levels, of which 73.89% (n=665/900) were unique to surface samples, 18.33% (n=165/900) to air while 7.56% (n=68/900) were shared between them (**Figure S2C**). Similarly to the differences in microbial composition between sample groups, Jaccard dissimilarity scores between samples revealed that air and surface samples represented two distinct groups (PERMANOVA *R*^2^ = 0.054; *p-value* < 0.009) according to the presence/absence of contigs containing genetic elements (**Figure S2D**).

Similarly to the genera distribution trends described above, the genus *Pseudomonas* represented the most prevalent taxa in the surface dataset, accounting for 25.87% (n=2101/8122) of the total read count (**Figure 2**). *Pseudomonas* reads were identified in 75% (n=27) of the surface samples containing contigs with relevant genetic elements, and 17.8% (n=5) of air samples containing genetic elements. The Bacilli class had the second highest number of read counts, making up 6.8% (n=554/8122) of the total reads with genetic elements. The majority were found on air samples (n=6), while only two surface samples, S2 (ward E, SR 16, Bathroom Vent) and S21 (ward E, Clean Utility, Vent 2), contained Bacilli reads with relevant genetic elements. This was followed by the order Enterobacterales, which had the third highest number of read counts, accounting for 5.5% (n=452/8122) of total read count. These were found in both air (n= 11) and surface (n=9) samples across multiple wards. By comparison, the *Rahnella* genus was detected mostly in the air samples, with species *Rahnella bonaserana* (n=148), *Rahnella aceris* (n =89), and *Rahnella Victoriana* (n=57) accounting for most of the *Rahnella* reads with relevant genetic elements in the air microbiome (**Figure 2**).

**Figure 2.**
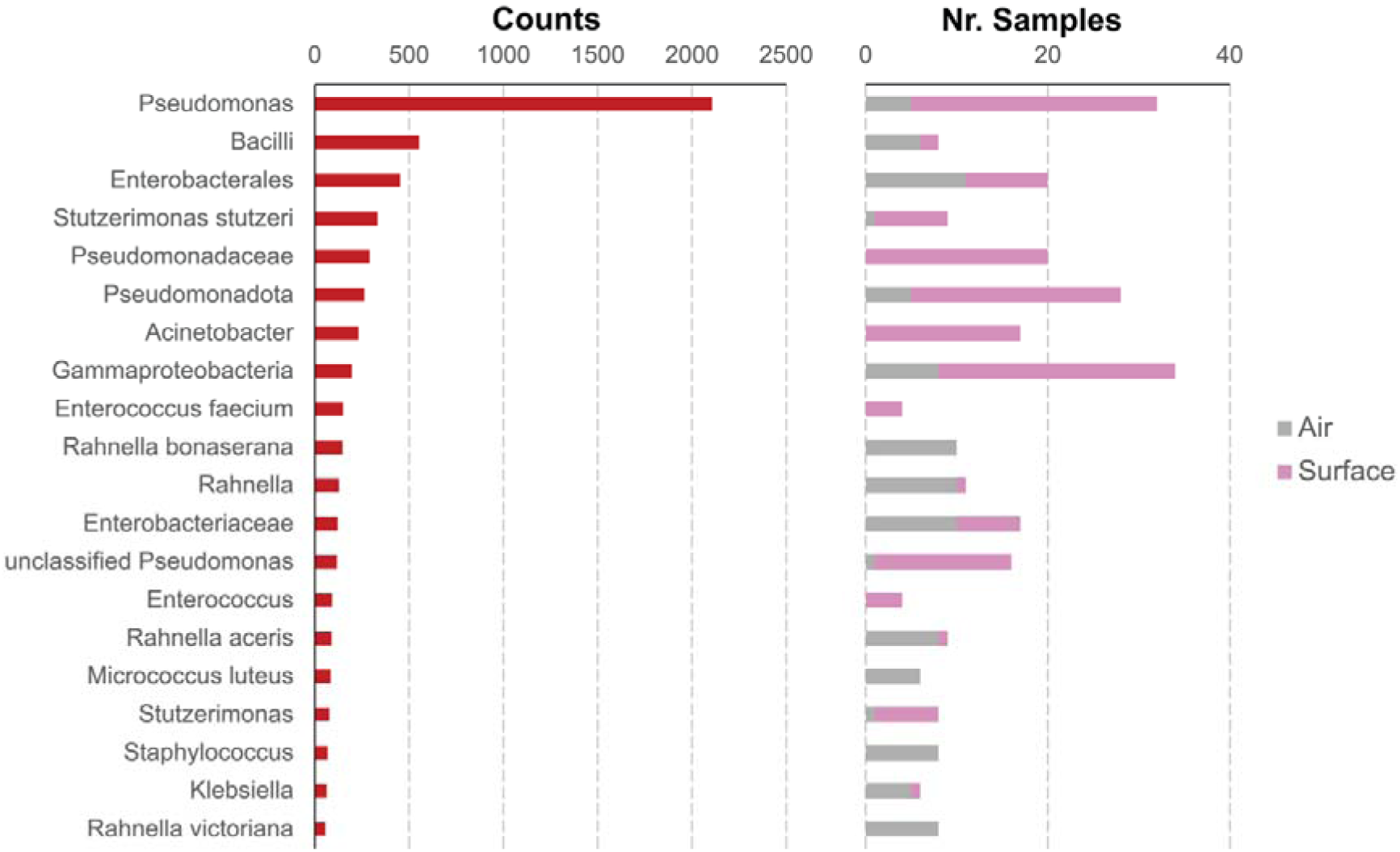
Barplots showing the number of contigs (counts) for the top 20 kraken2-annotated taxa with the highest number of contigs with relevant genetic elements, as well as their distribution across air and surface samples.

### 3.2. Antibiotic Resistance Genes, Virulence Factors and Mobile Genetic Elements profile in the hospital environment

To gain a comprehensive understanding of the resistance mechanisms, pathogenesis, and mobility of the different taxa identified within the hospital environment, we conducted a detailed profiling of antimicrobial resistance genes (ARGs), virulence factors (VFs), and mobile genetic elements (MGEs) (**Figure 3**).

**Figure 3.**
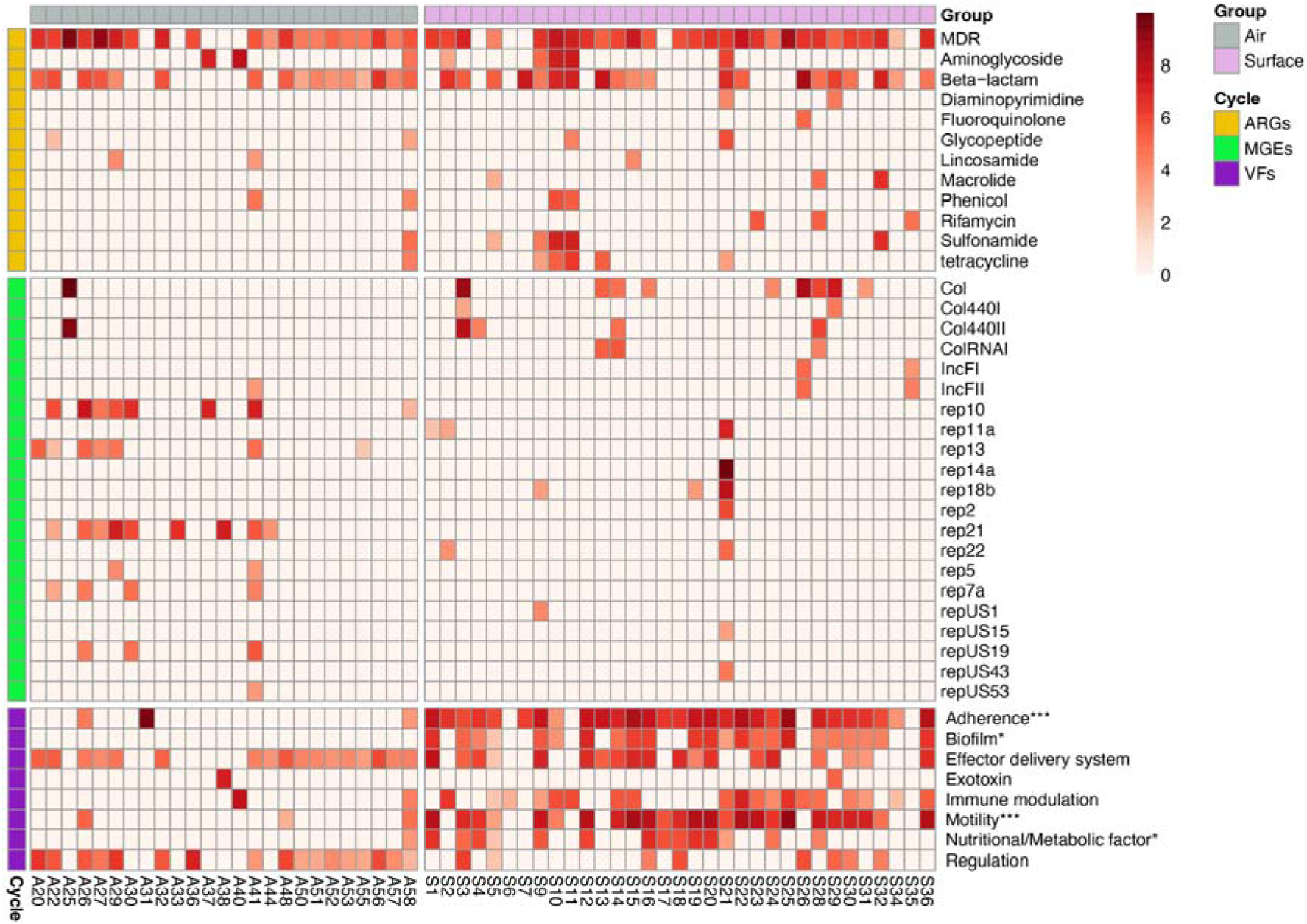
Heatmap showing the prevalence (presence/absence) and abundance (in rpm) of the main types and pathways of ARGs, MGEs, and VFs, across the sample set. Pathways/types of genetic elements that were calculated to be differentially prevalent and abundant across the two sample groups (air vs surface) using the Maaslin3 test are highlighted using the conventional nomenclature for significance (*p-value* < 0.05 (*); *p-value* < 0.01 (**); *p-value* < 0.001 (***)).

A total of 1964 reads were identified to contain antibiotic resistance genes (ARGs), and 123 unique ARGs were identified across different sample groups at the hospital (**Figure S3A**). Genes involved in multidrug resistance (MDR) and resistance against beta-lactams were found to be the most prevalent and abundant ARGs across the dataset, being detected in 48 (20 air/28 surface) and 36 (16 air/20 surface) samples, and with an average of 1024 and 494 reads per million (rpm), respectively (**Figure 3**). Notably, the multidrug resistance (MDR) gene *H-NS*, which acts as a histone-like protein involved in global gene regulation of Gram-negative bacteria, was the most common gene found in the dataset (n=265) (**Figure S3B**), and mostly represented in air samples across multiple wards (**Figure S4**). This MDR gene was identified in reads classified as various taxa within the orders *Enterobacterales*, *Vibrionales*, and *Pseudomonadales*, including the genus *Rahnella*. Similarly, the *CRP* gene, the second most abundant ARG (n=220) in the dataset, was found predominantly in air samples, and was primarily associated with reads classified as species from the *Rahnella* genus described earlier. Several other genes associated with MDR were found mainly in surface samples. These include the gene *yajC* (n=133), which was found across multiple reads belonging to the genus *Pseudomonas* and *Stutzerimonas* including the species *Stutzerimonas stutzeri*. The resistance genes *mexF, mexK, and mexB,* which are all part of the *Mex (MexAB-OprM)* efflux system, were also identified primarily in surface samples across several wards (**Figure S4**) and accounted for 5.8% (n=144/2462) of the total ARGs in the dataset. Of clinical importance was the detection of the wide-spectrum *bla_OXA_* type genes, which confer resistance to carbapenem, cephalosporin and penam, in surface samples across different wards and air sample 58 (ward G, SDEC Corridor) (**Figure S4**). Many subtypes were present such as *bla_OXA-10_*, *bla_OXA-284_*, *bla_OXA-134_*, *bla_OXA-647_*, and *bla_OXA-648_*accounting for 10.03% (n=247/2462) of the total ARGs identified. Despite these differences in prevalence of specific ARGs, the Maaslin3 analysis of both prevalence and abundance of genetic elements revealed no significant differences in ARG types between the air and surface groups (**Table S2**).

**Figure S3.**
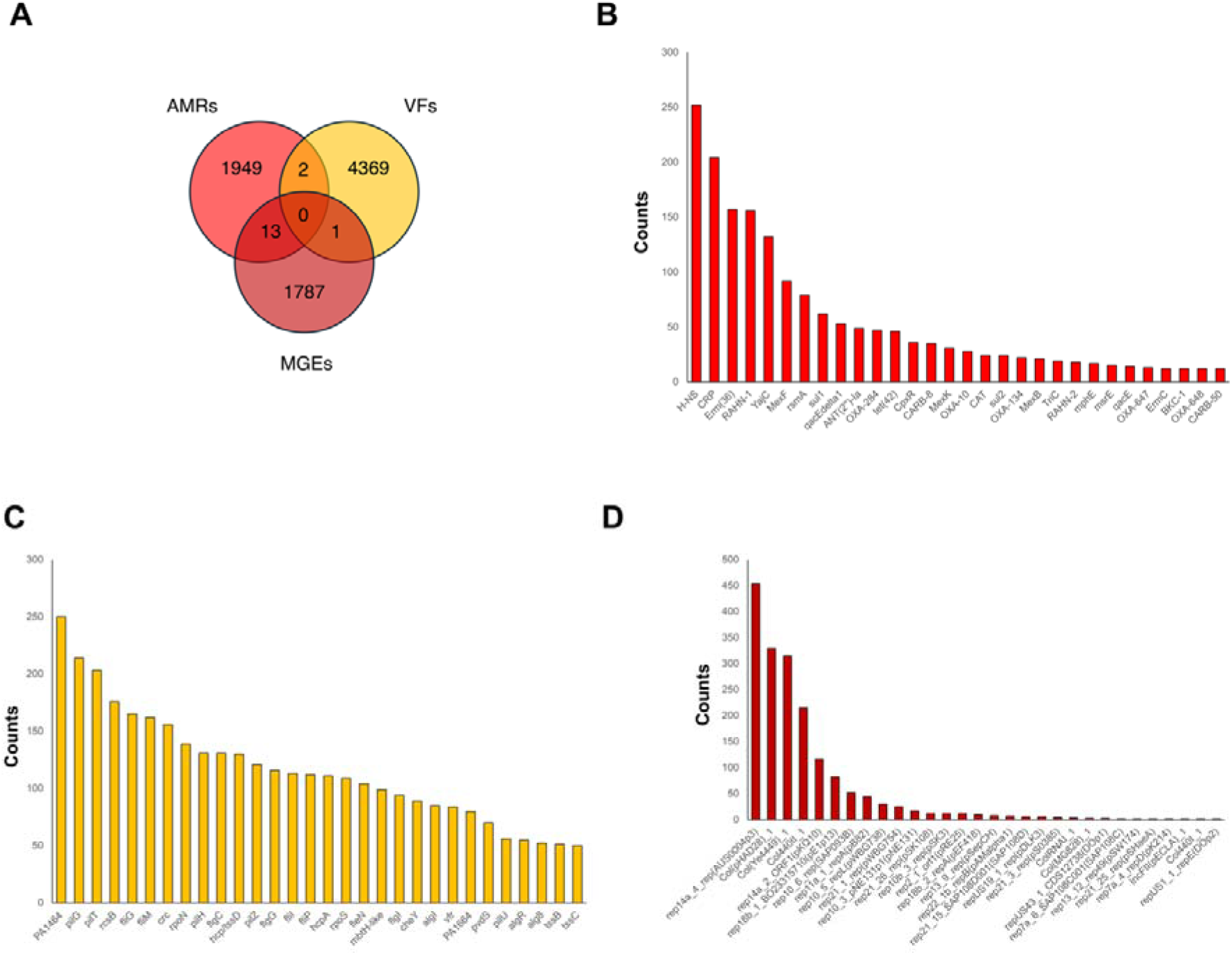
Number of reads with AMR, VFs, and MGEs detected in the dataset (**A**), as well as the counts for the top 30 most abundant genes for AMR (**B**), VFs (**C**), and MGEs (**D**) identified across the dataset.

**Figure S4.**
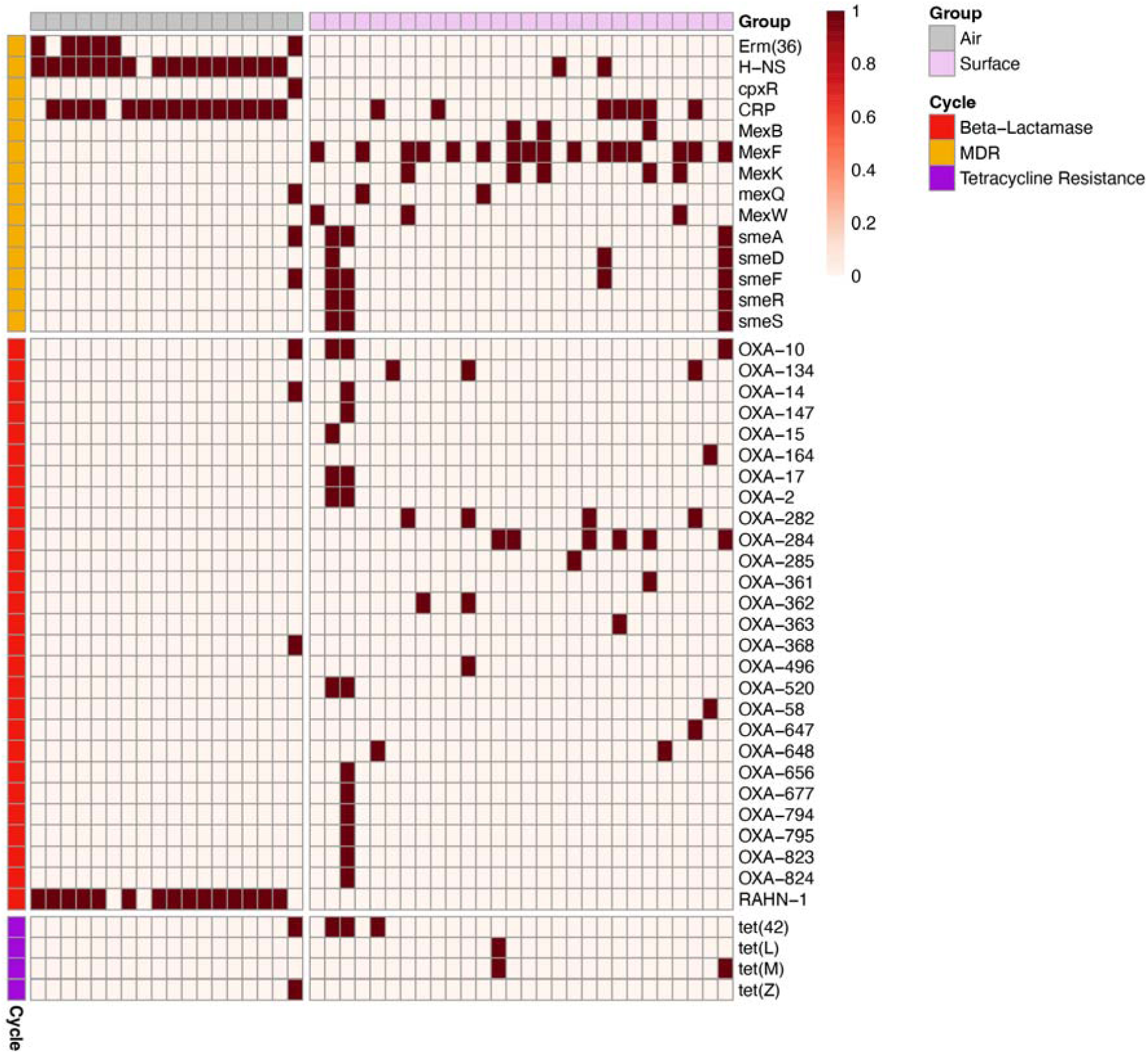
Heatmap showing the presence/absence of reads containing selected AMR genes across the sample set. Presence is denoted by the dark red cells on the heatmap.

A total of 4372 of reads contained virulence factor genes (VFs), with 106 distinct VFs being identified among various sample groups at the hospital (**Figure S3A**). Four distinct VF pathways were found to be significantly enriched (p-value < 0.05) in the surface samples, namely biofilm formation, adherence, motility, and nutritional/metabolic factors (**Figure 3; Table S2**).

Several type of secretion systems were detected across the dataset. For instance, the virulence genes *pilG, pilH, pilI, pilJ, pilM, pilR, pilT, pilU, pilX,* and *pilZ,* which are all associated with the type IV pili system, counted for 19.29% (n=1064/5496) of the total VFs (**Figure S3C**). Most of them were found in reads from surface samples across multiple wards, except for 16 counts which were found in the air samples 26 (ward C, Bed bay) and 58 (ward G, SDEC Corridor). Similarly, the type VI secretion system (T6SS) genes *hcpA*, *hcp1*, *hcp/tssD*, *tssB*, *tssC*, *hsiB1/vipA/tssB*, *hsiC1/vipB/tssC*, *icmH/tssL*, *tssG*, *tssF*, *tssE*, and *tssK* accounted for 12.56% (n=690/5496) of the total VFs count and were found in both air samples across multiple wards in the hospital. Notably, the gene *hcp/tssD* was prevalent in air samples and was associated with taxa belonging to the order *Enterobacterales*, including the genera *Enterobacter*, *Rahnella*, *Yersinia*, and *Buttiauxella*. In addition, biofilm-associated genes such as *algI, algR, alg8, algU, and algA* were also predominant in reads from several surface samples across multiple wards and accounted for 5.45% (n=300/5496) of the total VFs counts. They were also identified in reads belonging to various species from the *Pseudomonas* and *Stutzerimonas* genera including the pathogen *S. stutzeri.* Of note is also the detection of the *vfr* gene the hospital microbiomes, which encodes the virulence factor regulator (Vfr), a global regulatory protein that plays a crucial role in managing the expression of various virulence factors, including exotoxins, proteases, and and biofilm formation^48^. This *gene* accounted for 1.53% (n=84/5496) of the VFs counted and was predominantly found in reads from surface samples collected from multiple wards of the hospital, which were associated with various species from the *Pseudomonas* and *Stutzerimonas* genera, including the pathogen *S. stutzeri*. On the other hand, the *pvdS* gene, which encodes an alternative sigma factor known as pyoverdine (PvdS) vital for regulating virulence factors and iron acquisition^49^, accounted for 1.30% (n=72/11055) of the total VFs. Like the *vfr* virulent gene, *pvdS* was mainly found on reads from surface samples across several hospital wards and was linked to various species within the *Pseudomonas* genus.

By comparison, a total 50 distinct MGEs were detected in 1801 reads across 30 samples (**Figure S3A**), with an almost equal split between air (n=14) and surface (n=16) samples (**Figure 3**). Notably, the plasmid replicons *Col(pHAD28)_1,* Col440II_1 and *Col(Ye4449)_1*, which frequently carry genes that confer resistance to antibiotics, accounted for 22.56% (n=502/2181), 16.35% (n=350/2181) and 16.28% (n=341/2181) of the total MGEs gene count correspondingly (**Figure S3D**), although none of these were found in contigs with other relevant genetic elements. They were also found predominantly in surface sample S3 (ward E, SR 16, Bathroom Vent), S26 (ward E, Staff room freezer, interior drain), and A25 (ward C, Dirty Room), suggesting these sites are hotspots for such plasmids. By comparison, the replicons *rep10_6_rep(SAP093B), rep10_5_repL(pWBG738), rep10_3_pNE131p1(pNE131), rep21_1_rep(pWBG754), rep10b_3_rep(pSK3), rep21_26_rep(pSK108), and repUS19_1_rep(pDLK3)* accounted for *2*.75% (n=60), 2% (n=43), 1.20% (n=26), 1.11% (n=24), <1% (n=16), <1% (n=12), and <1% (n=14) of the total MGEs and were present in air samples across multiple wards, indicating a widespread distribution in the hospital environment. No statistically significant differences were detected in the abundance or prevalence of MGEs between the two sample groups (**Table S2**).

It is worth pointing out that several sampling sites harboured multiple types of genetic elements, highlighting them as possible hotspots of AMR (**Figure 3**). Specifically, 23 samples (9 air and 14 surface) harboured reads with genes for at least one type in each genetic element class (ARGs, MGEs, and VFs). The samples S21 (ward E, Clean Utility, Vent 2), A41(ward K, Bed Bay 17-21), and A58 (ward G, SDEC Corridor) harboured the highest number of genetic element types (18, 14, and 14 respectively), with the former two containing a high prevalence and abundance of replicon types in addition to ARGs and MGEs. Other noteworthy sampling sites included S3 (ward E; SR 16, Fridge inside drain), which harboured a high abundance of Col replicons as well as reads with MDR genes, as well as samples S10 and S11 (ward E, Dirty Utility), which exhibited a wide range and abundance of AMR types, despite not containing reads with replicon elements.

#### 3.2.1 Genetic elements associated with pathogenic species

Several AMR genes could be correlated to clinically relevant bacteria (**Figure 4**). For instance, the *bla_RAHN-1_* gene, which confers resistance to cephalosporins, was exclusively detected in air samples (**Figure S4**) and was mainly associated with reads from species belonging to the *Rahnella* genus including the pathogen *Rahnella aquatilis* described before. A similar trend was observed with the MDR gene *Erm(36)*, which was detected in air samples and linked to the pathogens *Kocuria palustris* and *Micrococcus luteus.* Furthermore, contigs classified as critical pathogens present in both air and surface across different wards like *Enterobacter hormaechei*, *Klebsiella pneumoniae*, *Escherichia coli*, *Pseudomonas aeruginosa*, and *Acinetobacter lwoffii* were related with the *bla_OXA_* genes subtypes described above. Similarly, the tetracycline resistance genes *tet(42), tet(L), and tet(U)* were found in critical pathogens like *Enterococcus faecium and S. aureus* in both air and surface samples. Likewise, the *CpxR* involved in the activation of expression of RND efflux pump *MexAB-OprM in P. aeruginosa* was found in reads classified as species belonging to the genus *Pseudomonas* on specific surface locations. The multidrug efflux complex *smeD*, *smeF*, *smeA*, *smeR*, and *smeS* counted for 3.96% of total reads and was related to species belonging to the genus *Stenotrophomonas* including the opportunistic pathogen *Stenotrophomonas maltophilia-*related to HAIs. As with the *bla_OXA_* type genes, these were found mainly in surface samples and air sample 58 (ward G, SDEC Corridor) (**Figure S4**).

**Figure 4.**
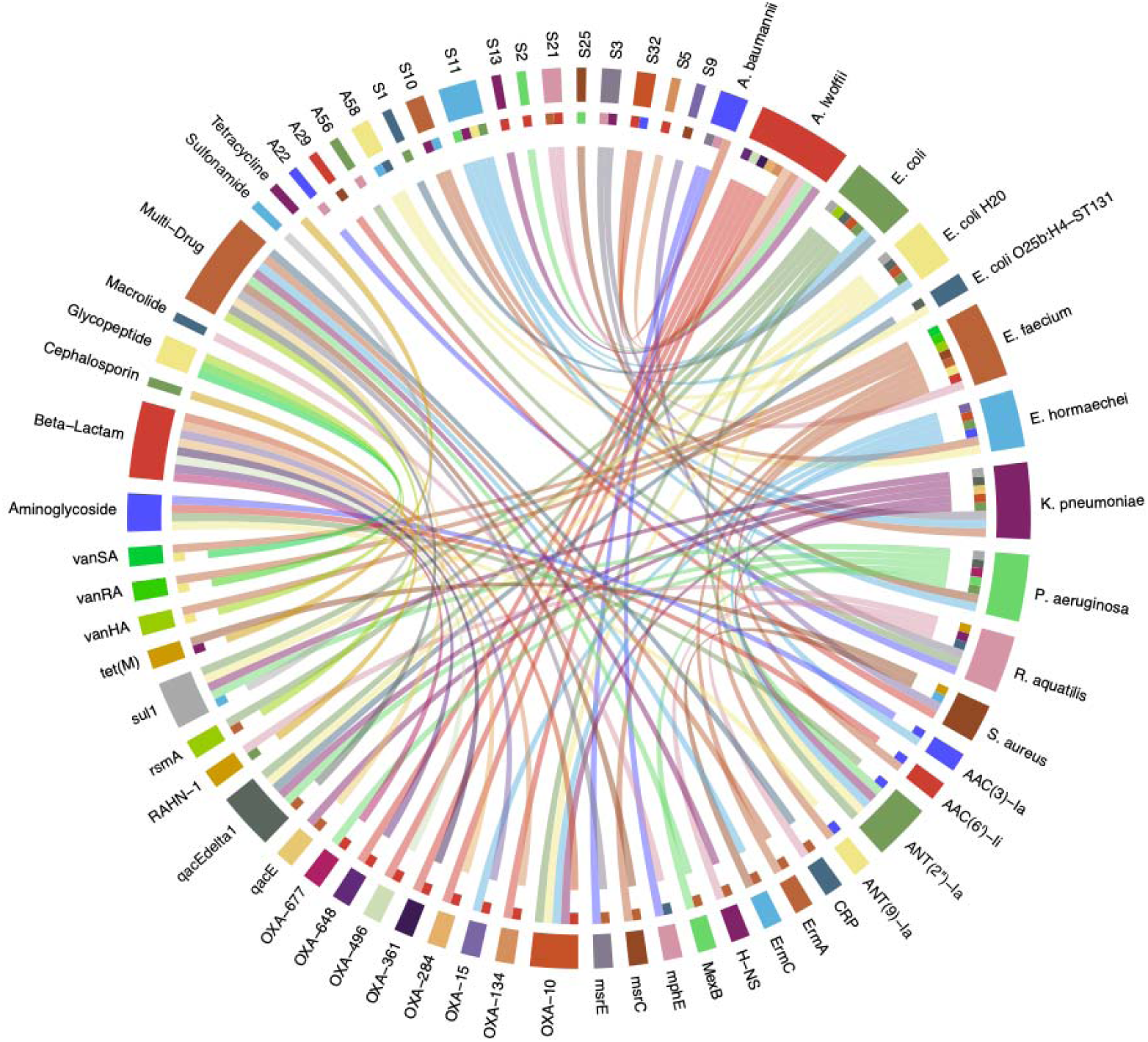
Chord diagram which highlights the clockwise relationship between the nodes: fifteen sample groups (Air and Surface), a selected group of nine critical and emerging pathogens, twenty-nine ARGs, and eight drug classes. The coloured strings represent the links between each node clockwise, starting from the sample node and ending on the drug class.

Similarly, several VFs and MGEs could also be mapped back to known pathogens. For instance, the majority of type IV pili system genes in the dataset were found in reads belonging to *Acinetobacter*, *Pseudomonas*, and *Stutzerimonas* genera, including pathogens such as *A. lwoffii*, *P. aeruginosa*, and *S. stutzeri.* Type II Secretion System (T2SS) genes such as *xcpT* gene wew also detected in reads belonging to genus *Pseudomonas* and *Stutzerimonas* including the pathogens *P. aeruginosa and S. stutzeri.* Notably, the gene *hcp/tssD*, which is part of the type VI secretion system (T6SS), was prevalent in air samples and was associated with taxa belonging to the order *Enterobacterales*, including the genera *Enterobacter*, *Rahnella*, *Yersinia*, and *Buttiauxella*. Among these, *Enterobacter hormaechei*, a well-known pathogen responsible for HAIs, was detected in sample 50 (ward I, Bed bay), 51 (ward I, corridor in the outpatient area) and 22 (ward B, Parents Kitchen). the plasmid replicons *Col(pHAD28)_1,* Col440II_1 and *Col(Ye4449)_1* were found in reads classified as various taxa within the orders *Enterobacterales*, including the pathogens *Enterobacter hormaechei, Escherichia coli, Klebsiella pneumoniae, Rahnella aquatilis,* and *Shigella flexneri*. Other pathogenic bacteria containing reads with replicons included *Enterococcus faecium, S. hominis*, *S. haemolyticus*, *S. epidermidis*, and *S. aureus* (**Table 1**).

**Table 1.**
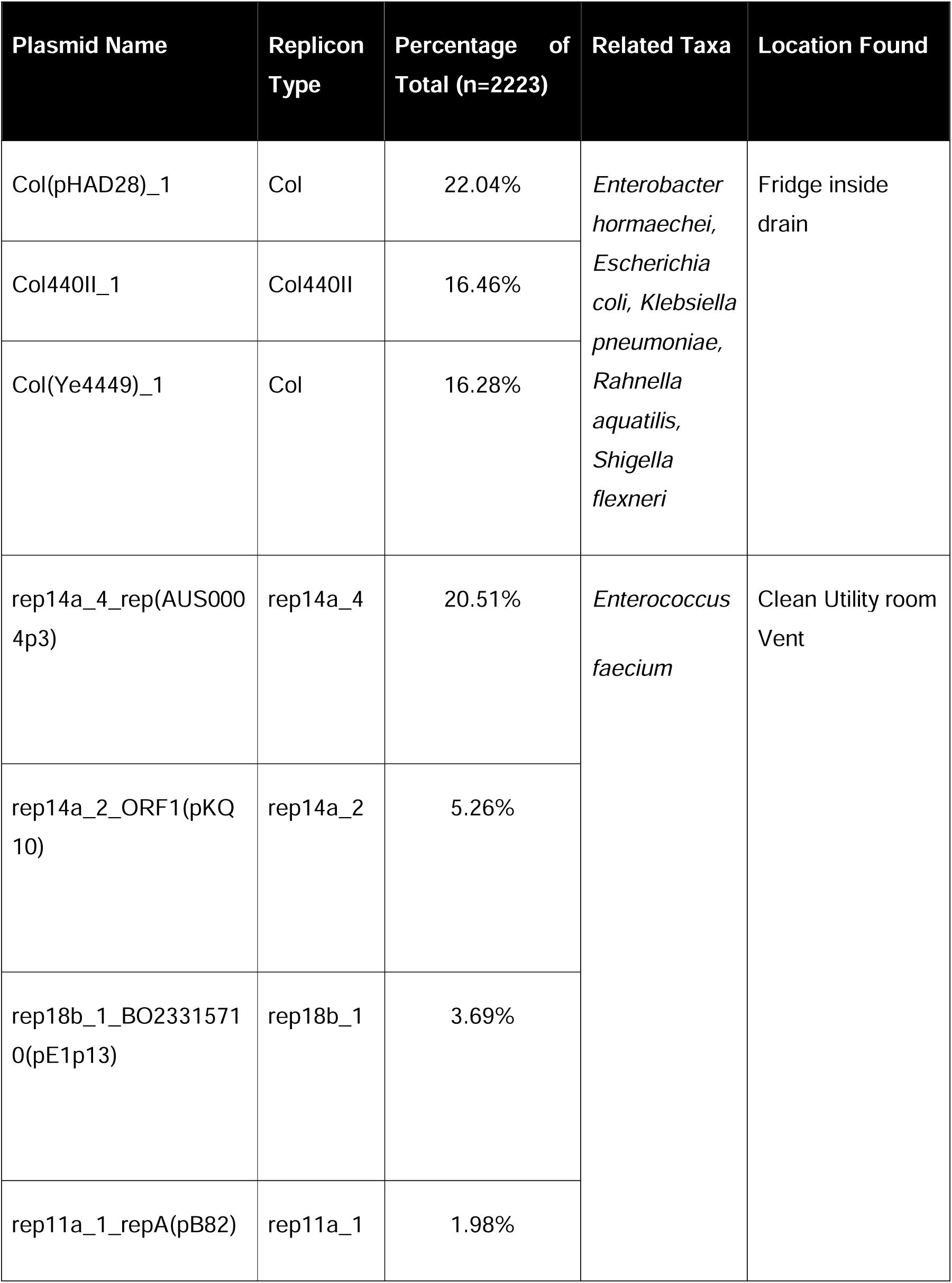
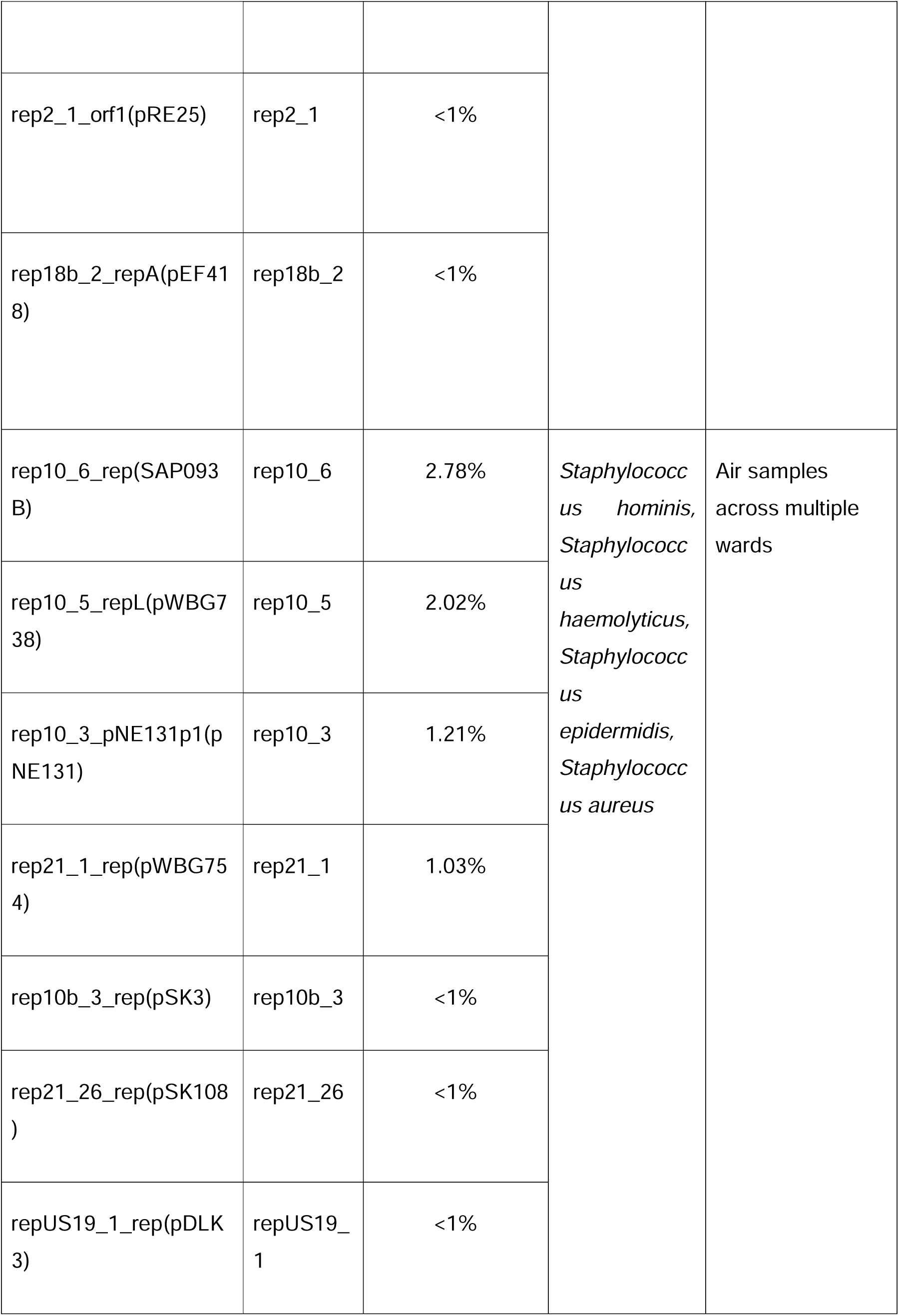
Summary of the top 16 Mobile Genetics Elements detected the hospital samples.

Despite the low number of reads (n=16) containing more than on distinct type of genetic element (**Figure S3A**), clustering reads by species revealed the potential co-existence of several distinct genetic elements within pathogenic bacteria (**Figure 5**). For instance, in *P. aeruginosa* several virulence factors were identified, including the Type VI secretion system (T6SS), the bacterial flagellum, and type IV pili, co-existed with resistance genes such as *bla_OXA-677_*, *sul1*, *qacEdelta1*, and *MexB*. Similarly, the *vanA* cluster, responsible for vancomycin resistance, *ANT(9)-Ia* and the *ErmA* gene (23S rRNA methyltransferase) co-existed with the virulence factors *acm,* encoding a surface adhesin protein in *E. faecium*, which plays a role in bacterial adherence to host tissues, contributing to virulence^50^ and *sgrA*, encoding a nidogen-binding LPXTG surface adhesin that is also involved in bacterial adhesion to host cells, enhancing the pathogen’s ability to colonise and cause infection^51^.

**Figure 5.**
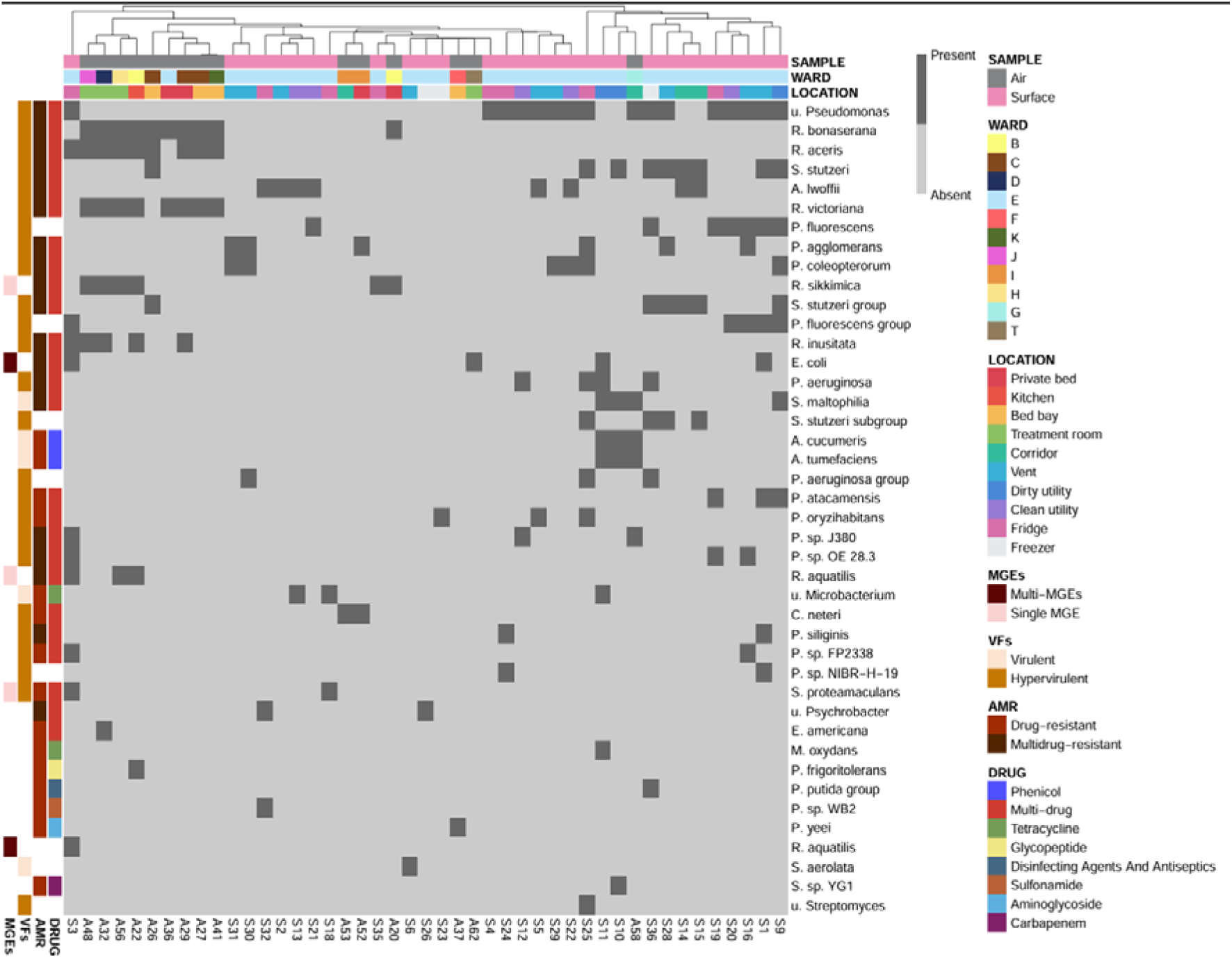
The figure represents a heatmap that provides a detailed visualization of the co-occurrence patterns among ARGs, VFs, and MGEs found in various taxa identified from the air and surface samples collected. In this visual representation, dark grey indicates the presence of specific taxa, while light grey signifies their absence. Each column on the left side corresponds to the various genetic elements present and their association with the taxa listed on the right, along with the relevant drug classes. The corresponding ward/location for each sample is also annotated on the top. Furthermore, the dendrogram offers a clustering of both sample groups based on the taxa present, highlighting similarities and differences in microbial composition across the hospital environment. The more representative taxa (n=42) at species level were plotted.

Multiple plasmid replicons co-existed with the above antimicrobial resistance and virulence factor genes in *E. faecium* described early. The *rep11a_1_repA(pB82)* plasmid was present which is reported for carrying the *repA* gene, often involved in the replication and stable maintenance of plasmids within bacterial cells. Their presence can facilitate the persistence and spread of AMR genes, such as vancomycin resistance, within hospital-associated *E. faecium* strains^52^. Another replicon present was *rep14a_2_ORF1(pKQ10)* which is associated with plasmids that can carry multiple resistance genes. The presence of ORF1 in these plasmids can enhance the bacteria’s ability to survive in environments with high antibiotic use, such as hospitals^52^. In addition, resistance genes such as *H-NS*, *CRP*, and *bla_RAHN-1_* co-existed with the transcriptional regulator *rcsB* and *hcp/tssD*, which is related to the Type VI secretion system. These genes were present in various species belonging to the genus *Rahnella*, including *R. aquatilis*, which was predominantly found in air sample (Figure 5). The plasmid replicon *Col(pHAD28)_1* was also present in *R. aquatilis* suggesting transfer mobility of these genetic elements within this emerging pathogen. A similar trend was observed in *S. aureus*, where *ErmC*, *ErmA*, *ANT(9)-Ia*, and *tet(U)* resistance genes, coexisted with various MGEs, such as *rep10_6_rep(SAP093B)*, *rep10_5_repL(pWBG738)*, *rep10_3_pNE131p1(pNE131)*, *rep21_1_rep(pWBG754)*, *rep10b_3_rep(pSK3)*, *rep21_26_rep(pSK108)*, and *repUS19_1_rep(pDLK3)*. Since these genetic elements were also found in other species of the genera *Staphylococcus, including S. hominis*, *S. haemolyticus*, *and S. epidermidis,* this result highlights the potential of HGT transfer within these species in the hospital environment.

In *K. pneumoniae*, resistance genes, including *ANT(2’’)-Ia (*aminoglycosides), *bla_OXA-10_ (*cephalosporins), *qacEdelta1 (*penams), *qacE (*antiseptics), and *sul1* (sulfonamides) coexisted with various MGEs, including *Col(pHAD28)_1*, *Col440II_1*, and *Col(Ye4449)_1*, which are known to confer antibiotic resistance or other advantageous traits to the bacterium. These genetic elements were also found in various taxa within the order *Enterobacterales*, including the pathogens *E. hormaechei, E. coli, R. aquatilis,* and *Shigella flexneri,* highlighting the transfer potential of HGT within common pathogens in the hospital environment. Furthermore, several of these GEs coexisted with various MGEs in *E. coli*, including *Col(MG828)_1*, *Col(pHAD28)_1*, *Col440II_1*, *and Col(Ye4449)_1*. In *E. hormaechei*, the resistance genes *bla_OXA-10_*, *ANT(2’’)-Ia*, *AAC(3)-Ia*, and *bla_OXA-15_* coexisted with *Col(pHAD28)_1* and *Col440II_1* plasmids replicon. In *A. lwoffii*, various *bla_OXA_* subtypes resistance genes such as *bla_OXA-134_*, *bla_OXA-282_*, *bla_OXA-284_*, *bla_OXA-361_*, *bla_OXA-496_*, and *bla_OXA-648_*coexisted with the virulence genes *pilG*, *pilT*, and *ompA*, which encode for twitching motility protein, flagellar basal-body rod protein, and outer membrane protein, respectively, and contribute to the bacterium’s motility and ability to establish infections.

### 3.4. Connectivity of the air and surface microbiome across hospital locations

Network analysis was performed to investigate how the reads with GEs identified in this study were connected across the hospital. This analysis revealed that several GEs belonging to clinically relevant bacterial species were widespread across both clinical and non-clinical locations within the hospital (**Figure 6**).

**Figure 6.**
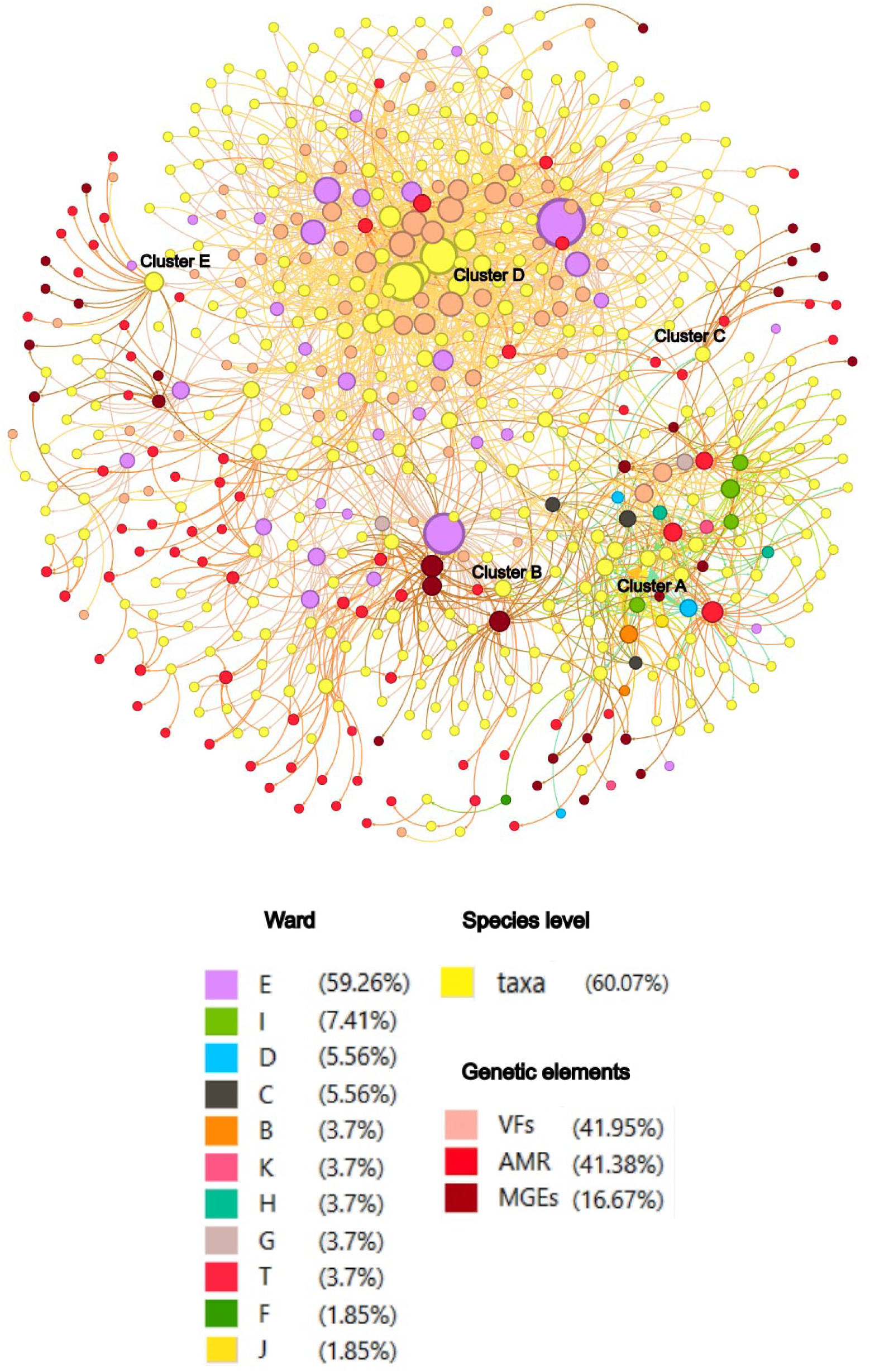
Network analysis and visualisation of the hospital microbiome cluster by air and surface samples. The network comprises various nodes representing different ecological components: sample groups (air and surface), bacterial taxa, antibiotic resistance genes (ARGs), virulence factors (VFs), and mobile genetic elements (MGEs). Node colours correspond to these categories (e.g., specific colours for ARGs, VFs, MGEs, taxa, sample groups). Edges indicate a direct association or presence (e.g., a sample contains a specific taxon, or a taxon contains a specific gene). The thickness of edges reflects abundance (taxa) and/or binary presence (genes). The Fruchterman-Reingold layout positions nodes with shared connections closer together, highlighting the composition of the air and surface microbiome in the hospital environment.

For instance, reads classified as taxa from the genus *Rahnella*, which was found to be the dominant genus in air samples, linked air and surface microbiomes across multiple wards (**Cluster A; Figure 6**). *R. aceris* was detected on a non!zlclinical surface (fridge drain, sample S3, ward E) and in air samples spanning diverse clinical and communal areas, including the Parents’ Kitchen (A22, ward B), patient beds (A26, A27, and A29, ward C), a treatment room (A32, ward D), a multi!zlbed bay (A41, ward K), an endoscopy rest area (A48, ward J), and the hospital gymnasium (A56, ward H). Similarly, reads belonging to *R. inusitata* were identified in overlapping air samples (A22, A29, A32, A48) and connected to the same surface sample (S3). Another highly abundant *Rahnella* species, *R. aquatilis,* was also detected in both air (A22, A56) and surface samples (S3). The reads for this species carried mobile genetic elements (Col!zltype plasmids). Lastly, *R. sikkimica* was detected across diverse hospital environments, including private beds (A20), treatment rooms (A32, A48, A56) kitchen (A22), and a fridge surface (S35), spanning five wards (B, D, J, H, E), suggesting environmental resilience and potential for cross-compartment persistence. Cluster A also highlighted the high connectivity of the *Rahnella* genus to ARGs (global gene regulators *HN-S* and *CRP,* plasmid-encoded elements *(RAHN-1*/*RAHN-2)* carrying a β!zllactamase (RAHN) conferring cephalosporin resistance), VFs (*hcp/tssD* (type VI secretion system), *rpoS* (RNA polymerase sigma factor) and *rcsB* (transcriptional regulator)) and MGEs (Col(pHAD28)_1, Col440I_1 and IncFII(pRSB107)_1) as previously described.

Another prominent cluster (Cluster B) was identified around *Enterobacter hormaechei*, a clinically significant member of the *Enterobacter cloacae* complex frequently associated with hospital!zlacquired infections. Unlike the environmental *Rahnella spp.*, *E. hormaechei* was widely distributed across both air and surface samples, underscoring its potential for persistence and dispersal within the hospital environment. Airborne detections included the Parents’ Kitchen (A22, ward B), Bay 50–53 (A50, ward I), Clinic E Corridor (A51, ward I), and the SDEC Corridor (A58, ward G). Surface reservoirs were equally diverse, spanning both clinical and non!zlclinical areas: Dirty Utility Vent 2 (S10, ward E), staff room freezer drain (S26, ward E), kitchen vents (S29, S31, ward E), and the fridge drain (S3, ward E). This clusters showed the co-occurrence of reads containing with resistance determinants (aminoglycoside!zlresistant *ANT(2’’)-Ia* and *AAC(3)-Ia*, and β!zllactamase genes including *bla_OXA_*_f*10*_ and *bla_OXA_*_f*15*_) VFs (hcp/tssD (type VI secretion system)), and MGEs (Col!zltype plasmid replicon, including *Col440II_1 CP023921* and *Col(pHAD28)_1 KU674895*).

A third large cluster (Cluster C) was formed by multiple *Staphylococcus* species. *Staphylococcus hominis* was widely distributed across wards, detected in air samples from the Parents’ Kitchen (A22, ward B), patient beds (A29, ward C; A30 and A33, ward D), a multi-bed bay (A37, ward F), and the Intervention CT suite (A44, ward K). Meanwhile, *S. haemolyticus* and *S. epidermidis* were both recovered from patient proximal air samples (A26, A27, A29, A41), while *S. aureus* was detected in overlapping air samples (A29, A30, A41) and uniquely on a surface reservoir (S9, Dirty Utility Vent 1, ward E). The network associated with *Staphylococcus* species revealed dense connections to resistance determinants and multi-plasmid replicons (**Table S3**). Among the *staphylococci* contigs with GEs identified in the dataset, those classified as *Staphylococcus aureus* emerged as the principal mobilome hub, carrying the broadest diversity of plasmid replicon families (rep10, rep10b, rep21, repUS19) in combination with clinically relevant resistance genes (*ErmC*, *tet(M)*). This contrasts with *S. hominis* and *S. epidermidis*, which were enriched for maintenance plasmids (rep21, rep7a) and antiseptic resistance (*qacG*), and *S. haemolyticus*, which was limited to a single rep10 replicon.

It is important to note that another cluster centred around *Pseudomonas spp.* reads (Cluster D) was notable for its dense virulence repertoire, combining motility, adhesion, biofilm formation, secretion systems, and global stress regulators. Its simultaneous detection in air and diverse surface reservoirs across wards G, C and E underscores its ecological versatility and persistence in hospital environments. Reads identified as *unclassified Pseudomonas* spanned both air (A58, SDEC Corridor, ward D) and multiple surface reservoirs across wards E, including, vents (S1, S5, S16, S28, S29), clean (S12, S20, S22) and dirty utilities (S9), fridges (S3, S4, S19, S24, S25), and freezers (S36), highlighting its persistence across both clinical and nonclinical environments. *S. stutzeri* was also detected in an air sample (A26, Bed Bay, ward C) and multiple surfaces (S1, S10, S14, S15, S25, S28, S36, S9) across wards E, including bed bays, vents, dirty utilities, corridors, fridges, and freezers. Other species, such as *Pseudomonas aeruginosa* and *P. oryzihabitans* showed a surface-restricted distribution (S1, S5, S12, S23, S25, S36), and two highly connected nodes S3 and S25, were dominated by multiple *Pseudomonas spp*.

*The Pseudomonas spp.* subnetwork displayed a hypervirulent nature with an extensive arsenal of virulence traits, including flagella, Type IV pili, alginate biosynthesis, and Type VI secretion systems; combined with multidrug efflux pumps (MexF, MexQ/MexW/MexK) and metabolic regulators (*crc*), alongside global regulators (*rpoN*, *rpoS*, *fleQ*, *fleN*, *vfr*, *pvdS*, *CpxR*) and accessory loci (e.g., siderophore biosynthesis, LPS modification, chemotaxis) further reinforced their adaptability and pathogenic potential. Importantly, *P. aeruginosa* and *P. oryzihabitans* carried additional clinically relevant ARGs (*sul1, ANT(2”)-Ia, OXA-677, qacEdelta1*), reinforcing their role as resistance reservoirs on hospital surfaces. This constellation of traits highlights the ecological persistence and hypervirulent nature of *Pseudomonas* in the hospital environment, capable of bridging air and surface reservoirs across wards. The cooccurrence of efflux- mediated ARGs with multiple virulence factors highlights this taxon as a potential opportunistic pathogen capable of both resisting antimicrobial pressure and establishing- colonization.

Lastly, the *Enterococcus faecium* subnetwork (Cluster E) was surface-restricted, across wards E, including SR vents (S1, S2), clean utilities (S21) and Beverage Bay Fridge (S19). *E. faecium*-associated cluster includes a diverse array of genetic elements: including vancomycin resistance *vanA* cluster components (*vanR*, *vanS*, *vanH*), macrolide (*ermA*, *msrC*) and aminoglycoside resistance genes (, *ANT(9)-Ia*, *AAC(6’)-Ii*), and virulence factors (*sgrA*-surface adhesion protein involved in biofilm formation and host colonization, and *acm*-collagen-binding adhesin, linked to tissue invasion and persistence in host environments), and plasmid replicons (rep11a, rep18b, rep22, rep14a, rep2, repUS15), collectively indicating a multidrug-resistant, mobile, and potentially hypervirulent genotype identified on hospital surfaces. The exclusive detection of this cluster on ward E surfaces suggests environmental persistence and potential for indirect transmission. Its absence from air samples may reflect localised contamination or surface-specific colonisation niches. The coexistence with multidrug resistance and hypervirulence traits raises concern for hospital-acquired infections (HAIs), especially in immunocompromised patients or those exposed to contaminated surfaces during care.

## 4. Discussion

Hospitals have long been recognised as hotspots for the emergence and transmission of MDR microorganisms^11^ and are increasingly regarded as high-risk environments for outbreaks of difficult-to-treat hospital-acquired infections (HAIs). Such infections pose a substantial threat to public health, particularly for vulnerable patient populations^53^. Consequently, there is a critical need for rapid and efficient surveillance approaches capable of capturing the genetic diversity circulating within the hospital environments, including that disseminated via the airborne microbiome.

In this study, we demonstrated that hospital environments represent significant reservoirs of AMR, as evidenced by the high diversity of pathogenic taxa and AMR-associated genetic determinants detected in the hospital surface microbiome. Furthermore, we provide evidence for the potential transfer of these genetic determinants between hospital wards via the air and surface microbiomes, highlighting a previously underappreciated pathway for intra-hospital dissemination and amplifying the risk of future outbreaks caused by AMR microorganisms.

The long-read metagenomic analysis of the air and surface microbiomes across several hospital wards revealed a large diversity of bacterial taxa, with a significant fraction (900 distinct taxa) harbouring AMR genes. The taxonomy results suggested a limited degree of cross-over between the air and surface samples. For instance, *Escherichia coli* (<1%, n=58/20,453), a pathogen frequently implicated in urinary tract infections (UTIs) and severe bloodstream infections^54^, was detected in both air and surface samples across various hospital wards. Similarly, *Staphylococcus aureus* (<1%, n=66/20,453), associated with a wide spectrum of infections ranging from mild to life-threatening^54^, was predominantly identified in air samples 29, 30, and 41 (all from patient rooms) and surface sample 9 (Dirty Utility Room Vent). *K. pneumoniae* (<1%, n=52/20,453), a pathogen responsible for pneumonia, bloodstream infections, wound or surgical site infections, and meningitis^55^, was primarily recovered from surface and air samples 3 (Fridge interior drain), 10 (Dirty Utility Room Vent), 11 (Macerator side port), 26 (Staff Room freezer interior drain), and 29 (Kitchen vent). This limited cross-over is not surprising, considering that a very small number of air samples were taken from the same ward (ward E) in which the surface samples were taken from.

Importantly, several bacterial species associated with HAIs were found to harbour multiple genes implicated in the emergence of AMR. These included *Pseudomonas aeruginosa* and *Pseudomonas oryzihabitans*, both well-characterised opportunistic pathogens known to cause HAIs, particularly in immunocompromised patients and those with invasive medical devices^54,56^. The clinical management of *Pseudomonas* infections has become challenging due to increased antibiotic resistance^54^. The findings of this study, showing the co-occurrence of multiple classes of AMR-associated genes within members of this genus, further highlight its risk as a potential reservoir for MDR. In addition, the fact that *Pseudomonas* signatures dominated the surface microbiome correlates well with the *P. aeruginosa* outbreak reported previously in the hospital sampled in this study. This observation further highlights the continued risk of recurrent outbreaks, particularly involving virulent *Pseudomonas* species harbouring multiple MDR determinants detected in this study.

While most infections are caused by *P. aeruginosa*, other Pseudomonadaceae may also give rise to severe infections, albeit less frequently. For example, *S. stutzeri* has been reported as a pathogen that may occasionally cause bacteraemia, respiratory infections, urinary tract infections, wound infections, endocarditis, and bone and joint infections^57^. While infections caused by *S. stutzeri* are relatively rare, they can occur in hospital settings, especially among patients with underlying health conditions or those who have undergone invasive procedures^57^. In our study, *S. stutzeri* was associated with several ARGs, VFs, and MGEs, also highlighting it as a potential risk organism in the hospital setting. The presence of numerous virulence factors in both *P. aeruginosa* and *S. stutzeri* is a cause for significant concern. These bacterial strains contained several hypervirulent marker genes, equipped with a diverse array of mechanisms that enable them to evade the host’s immune defences and facilitate the onset of infections. This complex arsenal not only enhances their ability to thrive in hostile environments but also poses a formidable challenge in clinical settings. In fact, previous studies have also reported the complex virulence systems amongst clinical strains of *P. aeruginosa* and *S. nitrititolerans* (a species within the genus *Stutzerimonas*), emphasising their ability to evade host immune defences and cause infections^58,59^. These bacteria were reported to possess numerous virulence genes, including those involved in toxin production, flagellum formation, and biofilm development, which contribute to their hypervirulent behaviour and adaptability in hostile environments^58,59^.

The order *Enterobacterales* includes several clinically important pathogens known to cause HAIs, such as *E. coli*, *K. pneumoniae*, *Enterobacter species*, *P. mirabilis*, and *S. marcescens*^60^. Hospital mortality and length of stay has been associated with *Enterobacterales* positive blood cultures, highlighting the detrimental impact of delayed antimicrobial susceptibility results and inadequate empiric antibacterial therapy on patient outcomes^61^. In this study, *E. coli, K. pneumoniae, E. hormaeche, R. aquatilis,* and *Shigella flexneri* harboured various members of *bla_OXA_* genes family, which is a main contributor to the increased burden of resistance to carbapenem, cephalosporin, and penam resistance in healthcare settings^62,63^. In addition, the Type VI secretion system (hcp/tssD) was also present in these species, potentially contributing to their pathogenicity. Additionally, plasmids belonging to the *ColE1*-like family were identified within this taxonomy group. Notably, these plasmids have undergone substantial functional transformation in recent decades, characterised by adecline in its traditional bacteriocin production functions and an increase in various antimicrobial resistance genes, particularly enzymatic factors, which include several extended-spectrum beta-lactamases and carbapenemases^64^. The shift in their plasmid cargo highlights their emerging role as key drivers in dissemination of antibiotic resistance. Consequently, plasmid-mediated spread of resistance genes among *Enterobacteriales* poses a significant risk for outbreaks that are increasingly difficult to control and treat^65^. Of particular concern, a *ColE1*-like plasmid encoding a type IV secretion system (T4SS) was recently identified in *Providencia rettgeri* and linked to a nosocomial outbreak in Mexico. The plasmid harboured resistance genes conferring resistance to both β-lactam antibiotics (*bla_NDM-1_*) and aminoglycosides (*aph(3’)-VI*), genetic features more commonly associated with *Acinetobacter* plasmids^66^. Our findings, together with the previous studies demonstrating the importance of *ColE1*-like plasmid in the dissemination of AMR, highlight the potential for horizontal transfer of resistance and virulence determinants across clinically relevant pathogens.

One such pathogen is *E. faecium*, which displays an inherent capacity to acquire and maintain resistance to antibiotics and to withstand diverse environmental stressors, enabling it to be implicated in a wide range of infections, including urinary tract infections (UTIs), bacteraemia, endocarditis, Intra-abdominal and pelvic infections, surgical site infections, and infections associated with catheters and other implanted medical devices^67^. In this study, vancomycin resistant *E. faecium* (VREfm) harboured multiple plasmid replicons, as well as a diverse repertoire of antimicrobial resistance and virulence genes. Notably, plasmid replicons such as rep11a_1_repA(pB82) and rep14a_2_ORF1(pKQ10) were identified, which may contribute to the persistence, adaptability and spread of these strains in hospital settings. This is of particular concern given that VREfm is responsible for substantial proportion of HAIs globally and is classified by the World Health Organization as a priority pathogen for which the development of new antimicrobial agents is urgently needed^68^. Therapeutic management of VREfm infections are challenging due to limited treatment options, and is associated with increased morbidity and mortality rates, especially in immunocompromised patients^69^. Our findings further highlight the ongoing and escalating threat posed by multidrug-resistant *E. faecium*in in healthcare environments.

Other emerging pathogens identified in this study also appear to act as potential reservoirs of AMR. Notably, in *R. aquatilis*, cephalosporins resistance genes were detected to co-exists with the transcriptional regulator *rcsB* and *hcp/tssD*, as well as with the Type VI secretion system (T6SS), and the plasmid replicon *Col(pHAD28)_1*. Similar findings have been observed in A. *baumannii*, where the T6SS genes contribute to antibiotic resistance and virulence^70^. The presence of regulatory virulence factors and a mobilizable plasmid backbone in *Rahnella* species highlights their potential role as a facilitator of gene exchange and emerging opportunist within hospital microbiomes. Furthermore, the widespread detection and relative abundance of *Rahnella- associated* AMR *gene* signals in air samples collected across multiple hospital wards suggest that these species can readily disseminate through the hospital airborne pathways. Given the emergence of *R. aquatilis* species in clinical settings and its ability to cause infections in vulnerable patient populations, including bacteraemia, sepsis, respiratory diseases, UTIs, and wound infections^71,72,73,74^, our findings highlight the need for enhanced surveillance to better characterise the epidemiology, transmission dynamics and clinical significance of spread of *Rahnella spp.* in hospital environments.

A similar pattern was observed in *S. aureus*, in which resistance genes, including *ErmC*, *ErmA*, *ANT(9)-Ia*, and *tet(U)*, conferring resistance to aminoglycoside and tetracycline, were found to coexisted with MGEs such as *rep10_6_rep*, *rep10_5_repL*. Notably, these genetic genes and MGEs were also detected in other *Staphylococcus* species, including *S. hominis*, *S. haemolyticus*, and *S. epidermidis*, supporting the potential for horizontal gene transfer in clinical settings. In addition, carbapenem-resistant *A. lwoffii* was found to harbour multiple virulence genes, including *pilG*, *pilT*, and *ompA*, which contribute to the bacterium’s motility and its ability to establish infections. *A. lwoffii* has been reported as a persistent bacterium with a demonstrated capacity to acquire antimicrobial resistance^75^, and the absence of plasmid replicons in these isolates suggests that alternative mechanisms such as HGT mediated by non-plasmid elements, may have contributed to the acquisition of these resistance and virulence determinants^76^.

One of the key outcomes of this study is the identification of potential environmental hotspots for both established and emerging pathogens within the hospital setting, as well as insights into their possible transmission pathways. Notably, specific clinical locations, including vents surface of clean utility rooms in ward E and air samples collected from a bed bay in ward K were associated with a high abundance of genetic determinants associated with AMR. These areas warrant further investigation to determine potential sources of contamination by pathogenic or virulent species and to inform refinement of existing cleaning, ventilation and sterilisation protocols. Several clinically relevant pathogens were detected across multiple locations and wards. For example, *S. aureus* was predominantly identified in air samples from patient rooms in wards C, D and K (samples (S) 29, 30, and 41), as well as in surface samples from a dirty utility room in ward E, indicating potential cross-ward transfer. *A. lwoffii* was primarily detected on surfaces, including vents in clean utility areas (S21), SR 16 room and bathroom vents (S1 and S2), and the beverage bay refrigerator (S19). *A. lwoffii* has been implicated in healthcare-associated infections, including bacteraemia, pneumonia (particularly ventilator-associated pneumonia), urinary tract infections, wound infections, and infections in immunocompromised patients, particularly in intensive care and neonatal units^77^. Similarly, *Rahnella spp*. were found across multiple clinical and non-clinical environments, including refrigerator surfaces in non-clinical areas and air samples from treatment rooms, multi!zlbed bays, an endoscopy rest area and the hospital gymnasium. Additional evidence of cross-environmental distribution was observed for *E. hormaechei*, which was detected in air samples from parents’ kitchens as well as multiple clinical locations across different wards of the hospital. Similarly, *P. aeruginosa* was identified on a variety of hospital surfaces, including dirty and clean utility rooms, a stem cell refrigerator, interior drains, and kitchen vents, findings that may be consistent with residual environmental contamination linked to a previously reported outbreak. The observed overlap between non-clinical and clinical facilities indicates plausible transmission routes for virulent and MRD associated genetic signals within the hospital. These findings suggest potential weaknesses in infection prevention and control practices with respect to limiting microbial exchange between clinical and non-clinical areas, highlighting the need for enhanced environmental surveillance and targeted intervention strategies.

Overall, the findings of this study demonstrate that the use of low-biomass optimised metagenomics pipelines can substantially enhance hospital surveillance efforts aimed at monitoring and mitigating the emergence and dissemination of hospital-acquired infections. By enabling the detection of established and emerging pathogens throughout the hospital environment, as well as associated AMR determinants, these metagenomics pipelines provide clinicians and infection control teams with actionable tools to rapidly identify potential environmental reservoirs of HAIs and characterise their resistance profiles. Such information can directly inform evidence-based infection prevention and control strategies and antimicrobial stewardship practices in healthcare settings^78^. Moreover, our metagenomics pipeline enables the integration of pathogen detection with the identification antimicrobial resistance genes, virulence factors, and mobile genetic elements, thereby offering deeper insights into the hospital microbiome as a potential reservoir and transmission network for AMR.

Nevertheless, shotgun sequencing has limitations when compared to culture-based approaches. Specifically, it mainly detects abundant taxa, provides indirect inference of resistance or pathogenic potential, and does lacks information on phenotypic characteristics. To address these constraints, future studies should integrate culture-based methods and isolate level whole genome sequencing to complement the metagenomics approach developed here. Such a multimodal approach would enable more robust validation of resistance phenotypes and pathogen viability, ultimately resulting in a more comprehensive understanding of the dynamics and risks associated with HAIs.

## 5. Conclusion

This study provided insights into the hospital microbiome using a rapid and efficient metagenomic pipeline, demonstrating its effectiveness for rapid detection of pathogens, genetic determinants associated with antimicrobial resistance and virulence, and comprehensive profiling of the broader microbiome, including unculturable and intrinsically resistant taxa. The detection of critical and emerging pathogens in both air and surface samples, many of which harbour various ARGs and VFs, indicates an increased risk of hospital-acquired infections. Importantly, while the identified high priority pathogens represent clear clinical concerns, a wide range of additional microbial taxa carrying these genetic determinants may also contribute to infection risk. These organisms may act as reservoirs for antimicrobial resistance genes and virulence factors, facilitating horizontal gene transfer and the potential emergence of novel multidrug-resistant and hypervirulent strains, thereby increasing the likelihood of treatment failure in HAIs. In summary, our findings highlight the need for enhanced infection prevention and environmental surveillance strategies, particularly to protect vulnerable patient populations at increased risk of adverse outcomes.

## Supplementary tables

**Table S1.** Metadata for each sample, including the ward and location.

**Table S2.** The MAasLin3 output for the classes of AMR genes that were found to be significantly distinct between groups in terms of prevalence and abundance.

**Table S3.** Resistance genes and plasmid replicons identified in *Staphylococcus spp.* in the air microbiome.

## Availability of data and material

The fasta/fastq files from the sequencing runs of the samples used in this study were deposited in NCBI, under the Bioproject PRJNA1402132, and will be made available to the public upon publication. The scripts used for the data analysis can be found on GitHub (https://github.com/Julio92-C/hosMicro/).

## Authors’ contribution

OP was responsible for the sample collection and optimization of the methodology for the sequencing and analysis of the low-biomass samples used in this study, and was a major contributor in the analysis of the data and writing of the manuscript. JCOC developed and validated the in-house bioinformatics pipeline used in this study, was responsible for the data curation and storage, and was a major contributor in data analysis and writing of the manuscript. PC was a major contributor in the development and optimization of the wet-lab methodology used in study and was involved in the revisions of the manuscript. PHL was involved in the validation of the bioinformatics pipeline used in the study and was a major contributor to the writing of the manuscript. SA provided access to the hospital sampled in this study, and was involved in the sample collection and revisions of the manuscript. DO was a major contributor in the development of the sampling methodology, and was involved in the sampling. RM was a major contributor in the conceptualization of the study’s experimental framework and sample collection strategy, and was involved in the writing and revisions of the manuscript. HM was the main contributor in the conceptualization of the study as well as development of the experimental framework used in the study. HM was also responsible for the funding of the project, and was involved in the writing and revisions of the manuscript.

## Human Ethics and Consent to Participate declarations

Not applicable, as no human samples were used in this study.

